# Human vault RNAs exhibit diverse expression patterns and inter-locus compensation

**DOI:** 10.64898/2026.06.17.732918

**Authors:** Wayne O. Hemphill, Arthur J. Zaug, Cody J. S. Hecht, Thomas R. Cech

**Affiliations:** Department of Biochemistry, University of Colorado Boulder; Howard Hughes Medical Institute and BioFrontiers Institute, University of Colorado Boulder

**Keywords:** vtRNA, noncoding RNA, paralogs, expression, compensation

## Abstract

In 1986, a class of small, noncoding RNAs was discovered in association with an enormous, enigmatic ribonucleoprotein complex – the “vault” particle – and thus dubbed vault RNAs (vtRNAs). However, it’s since been recognized that the vast majority (≥95%) of these noncoding vtRNA molecules are not associated with the mysterious vaults, raising questions about their potential independent function(s). Moreover, humans express four vtRNAs from two different loci, and debate has arisen about whether the vtRNA paralogs share any functional connection, and whether they should remain classified together. Herein, we report the expression patterns of the four *VTRNA* paralogs in a variety of human cell lines, including various single- and multi-gene *VTRNA*-knockout cell lines. Knockout of one or more of the three *VTRNA1* genes leads to increased expression of vtRNA2-1, suggesting that their biological functions are related. Additionally, we interrogated the effects of vtRNA knockout on HEK293T cell growth, viability, and viral infection, and were unable to replicate previously reported associations. Collectively, our findings point to a potential functional connection between even the most distantly related human vtRNA paralogs, while reaffirming that their biological roles and mechanisms still require critical study.

## INTRODUCTION

In 1986, Leonard Rome’s lab identified massive, ovoid bodies contaminating their rat liver coated vesicle preparations. Initial characterization revealed these particles to be ∼100,000 nm^3^ ribonucleoproteins (RNPs) – the largest RNP particles ever identified in eukaryotes – which they termed *vault* particles (Kedersha and Rome 1986). The vault RNP is composed of 78 units of major vault protein (MVP), a structural component that is both necessary and sufficient for vault particle assembly, ∼2 units of telomerase-associated protein 1 (TEP1) positioned at each “cap” of the vault structure, ∼8 units of a vault-specific polyADP-ribosylation enzyme (vPARP) positioned within the hollow vault particle, and multiple copies of a class of small, noncoding RNAs (ncRNAs) called vault RNAs (vtRNAs), which are suspected to localize around the vault caps and bind TEP1 (TANAKA and TSUKIHARA 2012).

In humans, four functional *VTRNA* genes have been identified – the *VTRNA1-1*, *1-2*, and *1-3* genes clustered together at one locus (*VTRNA1*) and the *VTRNA2-1* gene alone at another locus (*VTRNA2*), both on the long arm of chromosome 5 (Stadler et al. 2009; Gallo et al. 2022; van Zon et al. 2001). These *VTRNA* genes are transcribed by RNA Polymerase III (Pol. III) and produce transcripts of 88 nt (1-2 & 1-3), 98 nt (1-1), and 108 nt (2-1) (Stadler et al. 2009; Kickhoefer et al. 1993). In addition to their box A/B motif typical of type-2 internal Pol. III promoters, the *VTRNA* genes also exhibit unique external regulatory elements (Stadler et al. 2009; Vilalta et al. 1994; van Zon et al. 2001). Each vtRNA is thought to form a stem-loop structure with internal loops and bulges interrupting the base-paired region.

Over subsequent decades, the vault RNP has been implicated in multidrug resistance, cancer progression, viral infection and innate immunity, DNA damage response, and more, though many of these functional relationships are called into question by the largely phenotype-less MVP-knockout mouse models (Maniatis et al. 2025; Frascotti et al. 2021; Ma et al. 2024; Wang et al. 2020; Mossink et al. 2002, 2003). At present, much of the interest in vault particle function is related to its potential for storing/transporting molecular cargo (Maniatis et al. 2025; González-Álamos et al. 2024; Eichenmüller et al. 2003; van Zon et al. 2006). Regardless, the vault particle’s strong evolutionary conservation throughout eukaryotes and the recent identification of potential homologs in bacteria hint at a fundamental biological role for vaults (Santos et al. 2023; Sokolskyi 2019; Suprenant et al. 2007; Daly et al. 2013; Kedersha et al. 1990).

Since their initial identification in association with vaults, vtRNAs have transitioned into subjects of focused study, a fact largely attributable to their predominantly vault-independent existence *in vivo* (Gallo et al. 2022; Taube et al. 2024; Aghajani Mir 2023; Nandy et al. 2009; Kickhoefer et al. 2002). Much like the vaults themselves, vtRNAs are highly conserved throughout eukaryotes, yet mice lacking their only vtRNA (*Vaultrc5* knockout) appear phenotypically wild-type, leaving vtRNA function enigmatic (Stadler et al. 2009; Mosig and Stadler 2011; Prajapat et al. 2024).

The human vtRNAs have been implicated in various biological processes, including proliferation, tumorigenesis, drug resistance, apoptosis, autophagy, viral infection, and micro RNA production, but many findings are purportedly paralog-specific and have not necessarily been consistent between cell types or even independent, analogous studies (Taube et al. 2024; Aghajani Mir 2023; Gallo et al. 2022; Stok et al. 2025; Gallo et al. 2025). Thus, the exact functions of the vtRNAs remain an open question, though several compelling molecular mechanisms have been proposed for individual human vtRNAs. vtRNA1-1 has been proposed to directly bind p62 to negatively regulate autophagy (Büscher et al. 2022; Horos et al. 2019), while vtRNA2-1 has been proposed to directly bind and inhibit the double-stranded RNA activated protein kinase R (PKR) (Lee et al. 2011).

Herein, we establish the vtRNA paralogs’ copy numbers per cell and gene sequence conservation across multiple human cell lines. In addition, we develop *VTRNA1* single- and triple-knockout HEK293T cell lines to interrogate the human vtRNAs’ potential biological functions using growth, viability, and viral replication assays and RNA sequencing and to elucidate any inter-paralog coordination in their expression.

## RESULTS

### vtRNA paralogs have variable expression across human cell lines

The copy number of a ncRNA is of interest because it typically correlates with function. For example, RNAs required for translation or splicing are high copy number, whereas microRNAs (miRNAs), which have specific sets of targets, have lower copy number. Thus, we quantified the copies per cell (cpc) of each vtRNA paralog (Table 1) in HEK293T, HeLa, U2OS, MRT, iPSC, and BJ fibroblast human cells using quantitative reverse transcription PCR (RT-qPCR; Figure 1a) and northern blots (Figure 1b). It was not possible to probe for all four vtRNAs on the same northern blot due to size overlap, but two blots were sufficient to quantify all four. vtRNA1-1 and vtRNA1-3 exhibited relatively consistent expression across tested cell types, in the range of a few thousand cpc (Table 1). vtRNA2-1 had a more variable expression pattern, with little or no RNA detected in HEK293T and U2OS cells but about a thousand cpc in the other cell types (Table 1). These low copy numbers suggest that these vault RNAs may have a specialized rather than a general function (see Discussion).

**FIGURE 1.**
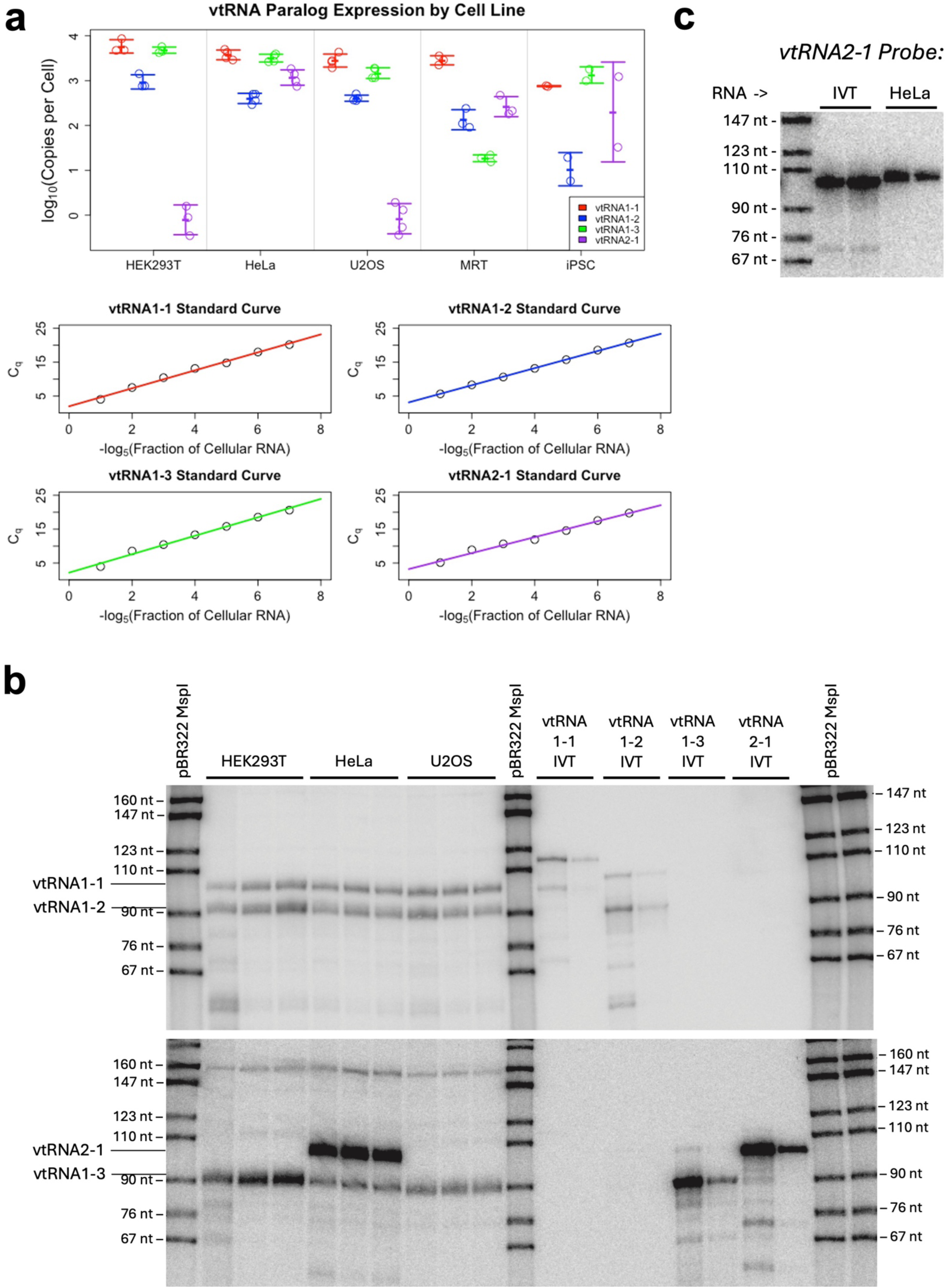
Common human cell lines have distinct vtRNA expression signatures. **[a]** Quantification of vtRNA1-1 (red), 1-2 (blue), 1-3 (green), and 2-1 (purple) copies per cell via RT-qPCR. Data points (top) indicate RNA samples from independent biological replicates, and error bars indicate mean ± standard deviation. Standard curves (bottom) are linear fits of known proportions of IVT-synthesized vtRNAs to total cellular RNA, versus their corresponding qPCR quantification cycle (C_q_) values. **[b]** Northern blots probing for vtRNA1-1 and 1-2 (top) and for vtRNA1-3 and vtRNA2-1 (bottom) in cellular RNA extracted from several human cell lines. Each group of three lanes represents three biological replicates. IVT RNA standards (“IVT”) with the full-length products indicated by the corresponding labels to the left of the gel. IVT also produced some transcripts larger or smaller than full length, which were included in the calculation of number of molecules. Copies per cell (cpc) for both northerns and RT-qPCR are reported in Table 1. **[c]** Northern blot comparison of the 100 nt vtRNA2-1 IVT product to the *in vivo* vtRNA2-1 (in HeLa cells).

**TABLE 1.**
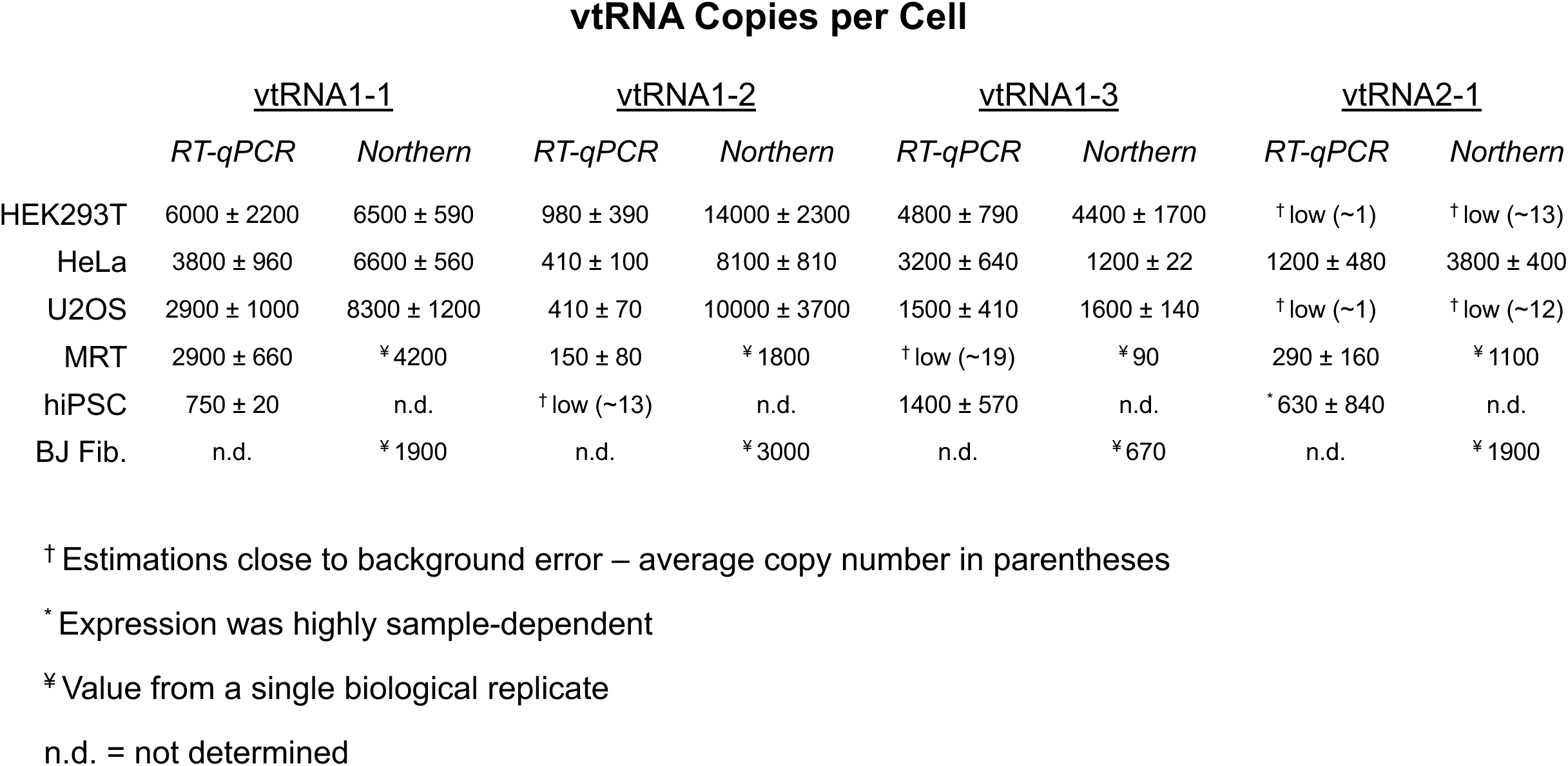
vtRNA paralog expression across human cell lines. Data indicate copies per cell (cpc) detected in each human cell line, reported as mean ± standard deviation across n=3 biological replicates. Values were calculated from Figure 1 RT-qPCR and Northern data using their respective standard curves.

In contrast, vtRNA1-2 exhibited substantially different cpc values between cell types. Perplexingly, its apparent levels also differed between methodologies, with northern blots averaging 10- to 20-fold higher vtRNA1-2 cpc values than RT-qPCR (Table 1). Yet, both methodologies used the same *in vitro* transcription (IVT) product for internal standard curves, the same purified RNA samples for input, and the northern blot probe was the RNA-complementary probe/primer from our RT-qPCR experiments. In addition, both methods appeared to specifically detect vtRNAs without notable off-target issues (Figure 1b and Supplemental Figure 1). Thus, while trends in relative vtRNA1-2 expression between tested cell lines are largely methodology-independent, the number of copies of vtRNA1-2 per cell as well as vtRNA1-2’s expression relative to other paralogs within a cell are less certain. By northern, vtRNA1-2 is often the most highly expressed paralog with up to ∼14,000 cpc, but by RT-qPCR, it’s the lowest expressed paralog, with several hundred cpc at most.

Given the variable paralog expression levels between human cell lines, especially for vtRNA2-1, we wondered if there might be cell type-specific differences in the vtRNA internal promoters or flanking sequences. Thus, we PCR-amplified the four human *VTRNA* genes from each tested cell type, sequenced the DNA, and aligned the sequences (Supplemental Figure 2). However, there was no sequence variation across tested cell types, either in the four *VTRNA* genes or within the ∼100 bp of upstream and downstream flanking sequences.

The vtRNA1-1 (RefSeq NR_026703.1), vtRNA1-2 (RefSeq NR_026704.1), and vtRNA1-3 (RefSeq NR_026705.1) transcripts are all annotated with a di-guanine start sequence and a poly-uracil terminal sequence common for Pol. III transcripts, yet the vtRNA2-1 (RefSeq NR_030583.3) transcript is annotated with eight “extra” nucleotides flanking these features (<u>CG</u>G_2_…U_4_<u>AUGCAA</u>). To determine whether this expanded sequence is representative of the vtRNA2-1 transcripts *in vivo*, we purified a vtRNA2-1 IVT control lacking these additional nucleotides and compared it to *in vivo* vtRNA2-1 by northern blot. Our results (Figure 1c) are consistent with vtRNA2-1 being transcribed (in our human cell types) as the annotated 108 nt RNA, affirming its sequence divergence from the other human vtRNA paralogs.

### Lack of *VTRNA2-1* expression in HEK293T cells appears epigenetically mediated

Given that all cell lines had the wild-type *VTRNA2-1* gene sequence, it seemed that the differences in vtRNA2-1 expression between cell lines must be attributable to epigenetic regulation or differentially expressed trans-acting factors (e.g., transcription factors). To test if the necessary trans-acting factors were available in HEK293T cells, we transfected them with a plasmid expressing *VTRNA2-1* under its own internal promoter (vector with *VTRNA2-1* plus 40 bp flanking sequences) and probed vtRNA paralog levels by northern blot. Our results (Supplemental Figure 3) revealed incredibly robust vtRNA2-1 ectopic expression, indicating that *VTRNA2-1* is suppressed epigenetically in wild-type HEK293T cells.

### *VTRNA* genes are efficiently deleted by CRISPR-Cas9

We used a CRISPR-Cas9 genetic editing approach (Figure 2a) to knock out vtRNA1-1 (vt1-1 KO; sites 1a/b), 1-2 (vt1-2 KO; sites 2a/b), 1-3 (vt1-3 KO; sites 3a/b), or all three *VTRNA1* paralogs (vt1-TKO; sites 1a/3b) in HEK293T cells, which are generally diploid for the *VTRNA* genes (Lin et al. 2014). We validated the knockouts by PCR (Figure 2), nanopore sequencing (Supplemental Figure 2 and 3a), northern blot (Supplemental Figure 4b), and RT-qPCR (Figure 3). CRISPR gene editing was so efficient at this locus that we obtained a preponderance of homozygous knockout clones. We were able to isolate one clone (CTRLdel) from the vt1-TKO genetic editing workflow that retained all of its *VTRNA* genes, which we hoped to use as an unedited control cell line. However, upon isolation of genomic DNA (gDNA) and DNA sequence analysis, it showed small deletions in the flanking sequences of *VTRNA1-1* and *1-3*, presumably due to CRISPR-Cas9 cleavage followed by non-homologous end-joining (NHEJ). Thus, the parental HEK293T cells were used as the control for comparison to the knockout clones. We found that the vt1-1 KO, vt1-2 KO, and two vt1-TKO (c1, c2) clones exhibited complete loss of their target vtRNAs, but the vt1-3 KO clone exhibited ∼2.5% residual vtRNA1-3 expression, and the vt1-TKO c3 clone exhibited ∼6.7% residual vtRNA1-2 expression (Supplemental Figure 4b and Figure 3).

**FIGURE 2.**
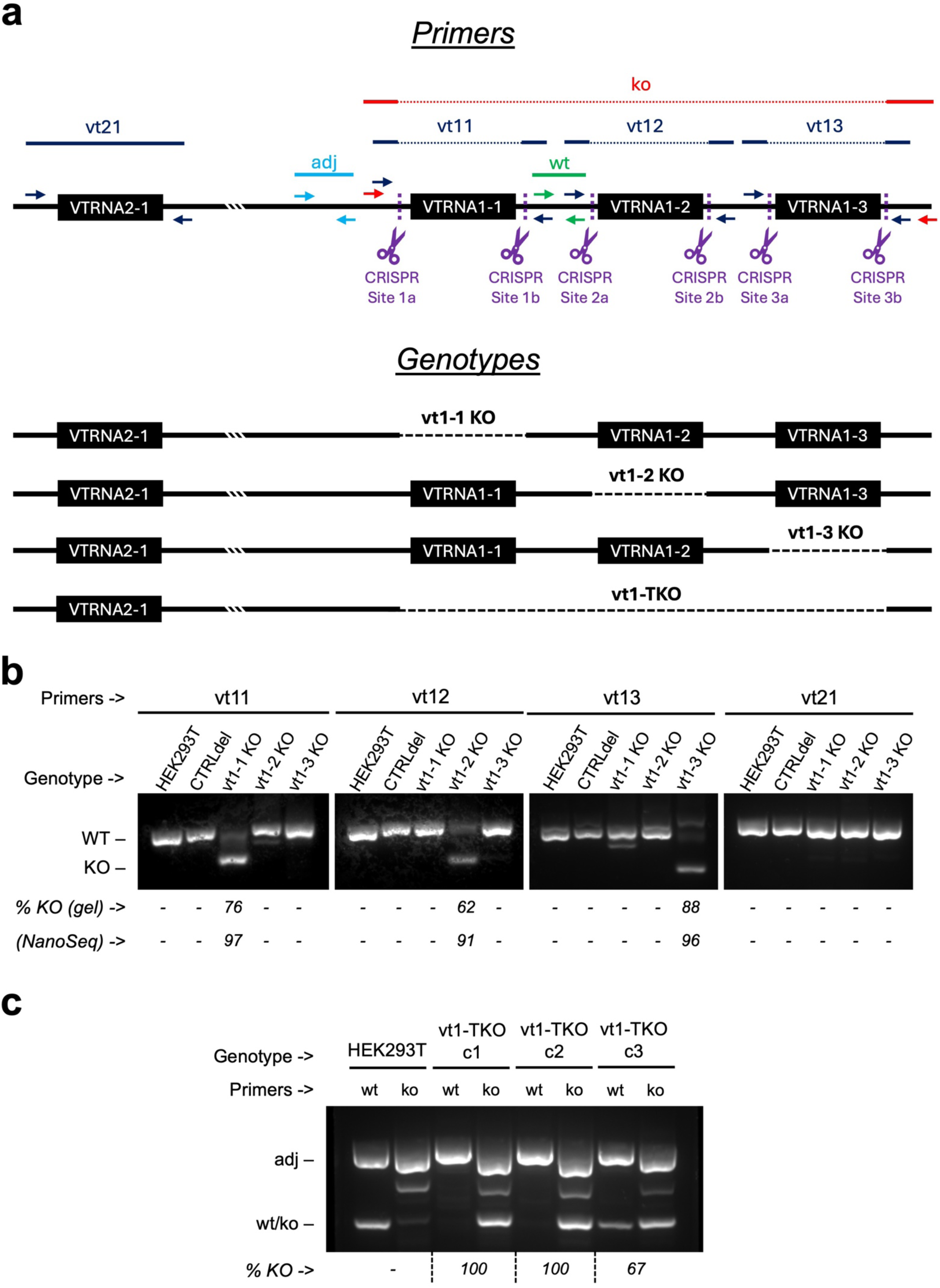
Knockout of vtRNA1 paralogs is non-lethal in HEK293T cells. **[a]** Schematics of the VTRNA1 and 2 genetic loci, CRISPR-Cas9 target sites (purple scissors) and vtRNA knockout genotypes (“Genotypes”), and PCR primer sets and their predicted amplification products (“Primers”). The triple white lines between the two VTRNA loci indicate their greater separation on Chromosome 5. **[b-c]** PCR was performed on gDNA from the indicated HEK293T cell lines (“Genotype”) with the indicated primer sets (“Primers”) from Figure 2a, and the products visualized by agarose gel electrophoresis. Efficiency of the single knockouts was quantified via densitometry (“% KO (gel)”) and nanopore sequencing of the PCR products (“NanoSeq”). The corresponding nanopore distributions for product lengths are in Supplemental Figure 4a, and the sequences of the dominant products are in Supplemental Figure 2. For the triple knockouts, the “wt” or “ko” primer sets were always run with the “adj” primer set as an internal control to allow normalization for densitometry-based calculations of knockout efficiency (“% KO”).

**FIGURE 3.**
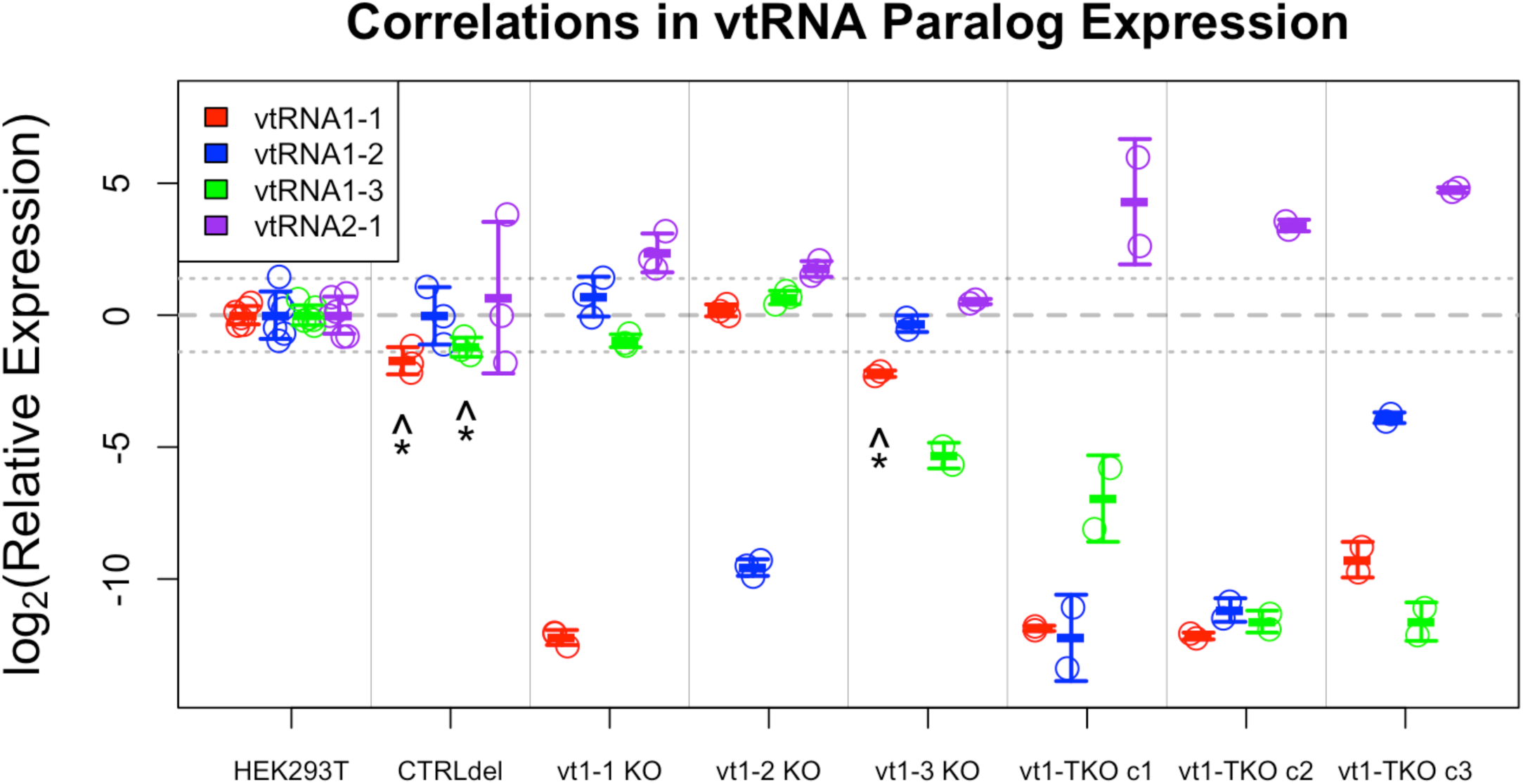
vtRNAs exhibit inter-locus but not intra-locus compensation. Quantification of vtRNA1-1 (red), 1-2 (blue), 1-3 (green), and 2-1 (purple) relative expression across HEK293T cell lines via RT-qPCR. Values correspond to ΔΔC_q_, which is calculated as vtRNA C_q_ normalized to GAPDH, then knockout ΔC_q_ normalized to wild-type (“HEK293T”) for each paralog. Data points are RNA samples from independent biological replicates, and error bars indicate mean ± standard deviation. Dashed grey line is a visual aid for no expression change, and dotted grey lines are a composite statistical threshold (p<0.05 in a two-sample t-test, assuming n=3 per group and variances equivalent to the “HEK293T” samples). Corresponding Northern blots are in Supplemental Figure 4. * Gene has a small, off-target deletion in its flanking sequence, identified in Supplemental Figure 2.

### *VTRNA* gene expression exhibits flanking sequence sensitivity

Early studies of vault RNA gene structures revealed the presence of unique, upstream regulatory elements in addition to their canonical, internal Pol. III promoter (Vilalta et al. 1994; van Zon et al. 2001). We identified several small, off-target deletions within our genetically edited cell lines (Supplemental Figure 2), which were in flanking sequences outside the vtRNA coding regions, yet always correlated with decreased vtRNA expression (Figure 3). In the vt1-3 KO cell line, a Δ_1_N_4_[VTRNA1-1] deletion (one missing base, four bases upstream of *VTRNA1-1*) correlated with a ∼5-fold decrease in vtRNA1-1 expression (Figure 3 and Supplemental Figure 2). In the CTRLdel cell line, Δ_4_N_5_[VTRNA1-1], Δ_4_N_5_[VTRNA1-3], and [VTRNA1-3]N_27_Δ_2_ deletions correlated with a ∼3-fold decrease in vtRNA1-1 expression and a ∼2-fold decrease in vtRNA1-3 expression (Figure 3 and Supplemental Figure 2). These deletions do not appear to directly impact the known *VTRNA1* external regulatory elements (TATA box, proximal sequence element (PSE), distal sequence elements (DSEs)) or DNA methylation sites, and are situated between the transcription start sites and TATA boxes, suggesting that the associated vtRNA expression defects are related to perturbed spacing of sequence elements that coordinate the transcriptional machinery (van Zon et al. 2001).

To test this, we transfected wild-type HEK293T cells with plasmids expressing vtRNA1-1 via its natural TATA box and internal promoter elements (vector with *VTRNA1-1* plus 40 bp flanking sequences), using either the wild-type gene fragment (Supplemental Figure 3, “VT11”) or a gene fragment containing the CTRLdel cell line’s Δ_4_N_5_[VTRNA1-1] deletion (Supplemental Figure 3, “VT11del”), then probed for vtRNA paralog levels by northern blot. Our results (Supplemental Figure 3) indicate that ectopic vtRNA1-1 expression is reduced with the CTRLdel versus wild-type gene fragment, affirming that the upstream deletions are responsible for altered vtRNA1-1 expression in our cell lines. In addition, we note that the transfected CTRLdel gene fragment appears to express a vtRNA1-1 molecule with erroneous length (Supplemental Figure 3), perhaps due to its altered spacing between the TATA box and normal transcription start site (TSS). However, this longer vtRNA1-1 was not visible in the CTRLdel cell lines, suggesting that the erroneous transcript may be an artifact of the missing PSE and DSEs in the ectopically expressed gene fragment.

Curiously, the off-target deletions upstream of *VTRNA1-1* also lie within an annotated enhancer (E1 hereafter; RefSeq LOC126807530) that overlaps most of the *VTRNA1-1* gene sequence, and which is annotated to affect expression of *VTRNA1-1* and 29 other genes. Similarly, the off-target deletion upstream of *VTRNA1-3* lies within a different annotated enhancer (E3 hereafter; RefSeq LOC112997562) that overlaps most of the *VTRNA1-3* gene sequence, and which is annotated to affect expression of *VTRNA1-2*, *VTRNA1-3*, and 19 other genes. However, neither the CTRLdel nor vt1-3 KO cell lines have perturbed vtRNA1-2 expression (Figure 3), despite their deletions in the E3 enhancer. Overall, these fortuitously generated mutations confirm that sequences upstream of the human *VTRNA1* genes have modulatory effects on their transcription.

### *VTRNA* gene expression exhibits inter-locus compensation

If the vtRNAs function in the same pathway, it seemed possible that the deletion of one paralog would be compensated by increased expression of another paralog. Considering our combined sequencing (Supplemental Figure 2) and expression (Figure 3) data for the single-knockout cell lines, there were no instances where the altered expression of a *VTRNA1* paralog could be attributed to the loss of another, suggesting there is no vtRNA intra-locus compensation in HEK293T cells. However, vtRNA2-1 was upregulated in the vt1-1 KO (∼5-fold), vt1-2 KO (∼3-fold), and all three vt1-TKO (11- to 27-fold) cell lines (Figure 3). Thus, it appears that the cells increase vtRNA2-1 expression to compensate for decreased levels of members of the vtRNA1 family. Importantly, there are no annotated enhancers affected by these gene deletions that are reported to regulate *VTRNA2-1*, implicating a novel inter-locus vtRNA compensation mechanism.

### HEK293T cells lacking vtRNA1 paralogs have normal growth and viability

In prior studies, human vtRNA1 paralogs have been associated with promoting cell proliferation and negatively regulating apoptosis, especially in cancerous cell lines (Amort et al. 2015; Taube et al. 2024; Bracher et al. 2020; Gallo et al. 2025, 2022; Aghajani Mir 2023; Hahne et al. 2021). We quantified relative proliferation rates of each of our vt1-TKO clones by co-culturing them with the parental WT HEK293T cells and using PCR with primers in the vtRNA1 locus to quantify any changes in the proportion of the two cell types over time (Figure 4a). We did not observe a significant (p = 0.114, two-tailed one-sample t-test) basal growth deficit in the triple-knockout cells. We measured apoptosis/viability in our single-knockout clones via flow cytometry with Annexin-V (membrane-impermeable phosphatidylserine binder) and propidium iodide (PI; membrane-impermeable DNA intercalator) staining (Figure 4b and Supplemental Figure 5). We did not observe an increase in basal apoptosis/viability in any of the single-knockout clones.

**FIGURE 4.**
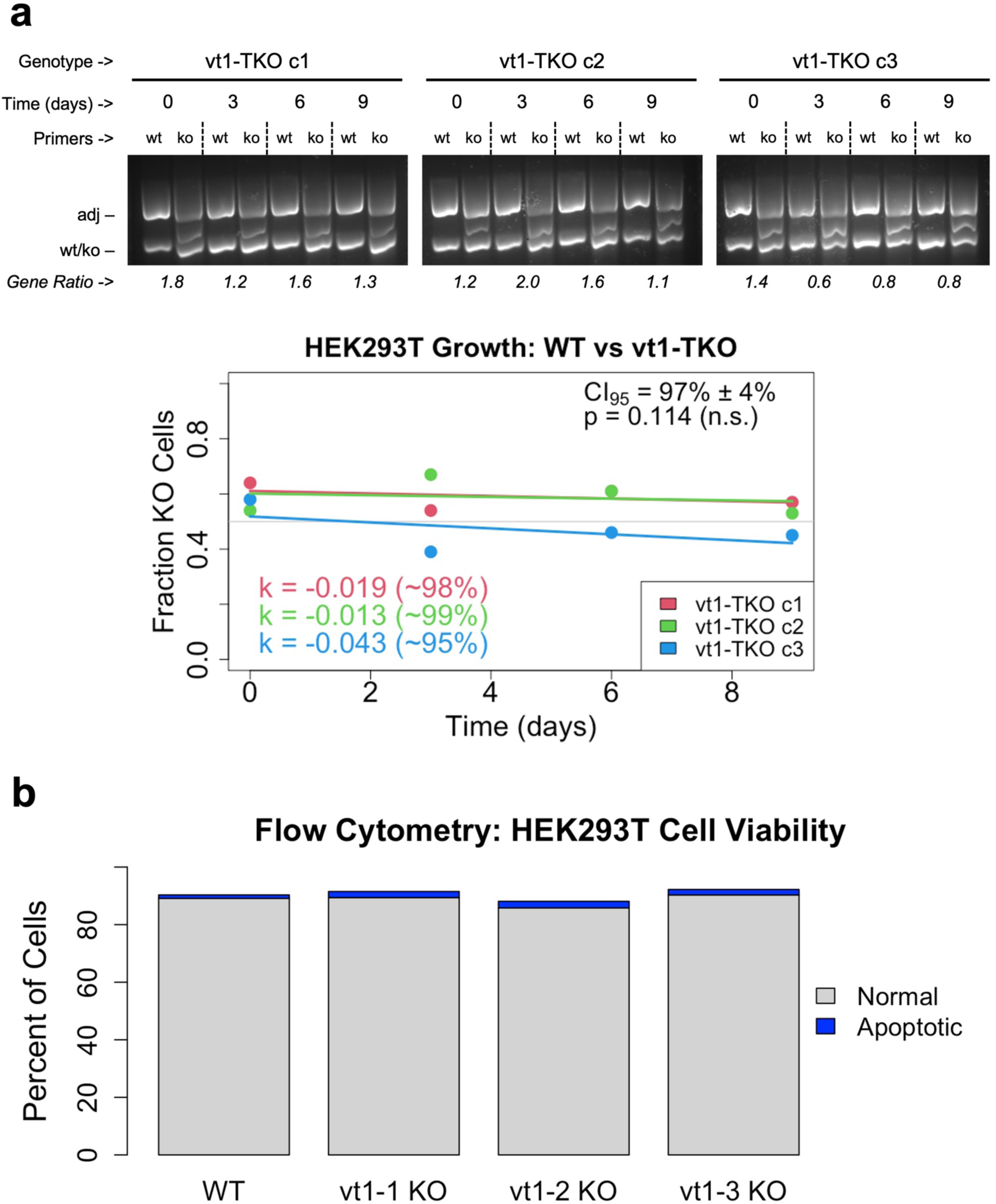
HEK293T cells lacking vtRNA1 paralogs exhibit no growth or viability deficits. **[a]** Mixed populations of wild-type (WT) and vt1-TKO HEK293T cells were grown for nine days with regular gDNA sampling, and their proportions tracked via PCR with indicated primer sets (“Primers”; see Figure 2a) and agarose gel electrophoresis. The prevalence of vt1-TKO versus WT cells (“Gene Ratio”) is the ratio of “ko” and “wt” product bands, each normalized to their respective “adj” product band. Plots of fraction KO cells versus time (bottom graph) for each triple-knockout clone were regressed (Eq. 1.1), and their rate constants (“k”, units of days^-1^) used to approximate relative growth rates for each clone (Eq. 1.2). 95% confidence interval (CI_95_) is from the clones’ relative growth rates, and p-value is from a one-sample t-test of the clones’ rate constants. **[b]** HEK293T cell lines were grown under standard conditions, dual stained with propidium iodide and Annexin-V, then analyzed via flow cytometry. Healthy (“normal”) and apoptotic cell percentages are taken from the corresponding gates in Supplemental Figure 5.

### vtRNA1 paralogs are not critical factors during OC43 viral infection

Several studies have implicated both the vault particles and vtRNAs in viral infection, particularly with enveloped RNA viruses (Stok et al. 2025; Liu et al. 2012; Wang et al. 2020; Ma et al. 2024; Peng et al. 2016; Nandy et al. 2009). Human coronavirus OC43 (HCoV-OC43) is an enveloped, single-stranded RNA virus – one of the seven known to infect humans, and one of the viruses associated with the common cold (Li 2016; Lau et al. 2011; Gaunt et al. 2010; Brüssow and Brüssow 2021). We measured relative OC43 replication in our vt1-TKO cell lines via an RT-qPCR-based assay (Supplemental Figure 6a) and relative vt1-TKO viability during OC43 infection via a PCR-based competitive growth assay (Supplemental Figure 6b). While one of our vt1-TKO clones exhibited increased susceptibility to OC43 infection/replication (Supplemental Figure 6a, vt1-TKO c3), the other two did not, suggesting the effect in clone 3 was not vtRNA-mediated. Similarly, we observed no significant difference in wild-type versus vt1-TKO HEK293T cell proliferation/survival during OC43 infection (Supplemental Figure 6b). Thus, at least for this virus in this human cell type, the three *VTRNA1* paralogs do not appreciably protect from or exaccerbate viral infection.

### vtRNA1 paralogs are not critical factors during RNA damage/repair

In bacteria, MVP homologs and ncRNAs sharing their operon have been implicated in RNA repair (Burroughs and Aravind 2016). To determine if human vtRNAs could play a similar role, we identified two strategies for cellular RNA damage: Schlafen family member 11 (SLFN11) overexpression and menadione (vitamin K_3_) treatment. SLFN11 is an endoribonuclease upregulated during apoptosis that cleaves a subset of tRNAs and leads to codon-specific ribosomal stalling (Elia et al. 2025; Iimori et al. 2025; Ogawa et al. 2025; Metzner et al. 2022). Menadione is a potent pro-oxidant that leads to broad cellular RNA damage, and which has heightened lethality against human cells lacking RtcB (RNA 2’,3’-cyclic phosphate and 5’-OH ligase) – a tRNA ligase known to share operons with MVP homologs in bacteria (Akiyama et al. 2022; Burroughs and Aravind 2016).

We quantified the effects of ectopic SLFN11 overexpression and repeated menadione treatment on survival/proliferation of our vt1-TKO (c2) cells by co-culturing them with the parental wild-type HEK293T cells and using PCR with primers in the vtRNA1 locus to quantify any changes in the proportion of the two cell types over time (Figure 5a). SLFN11 overexpression was confirmed by both RT-qPCR (Figure 5b) and western blot (Supplemental Figure 7). Menadione concentration was serially increased by ≤ 2-fold increments between treatments – from < 10% cell death to > 99% cell death (visually, via light microscopy) – to ensure that even modest biases in lethality would be captured.

**FIGURE 5.**
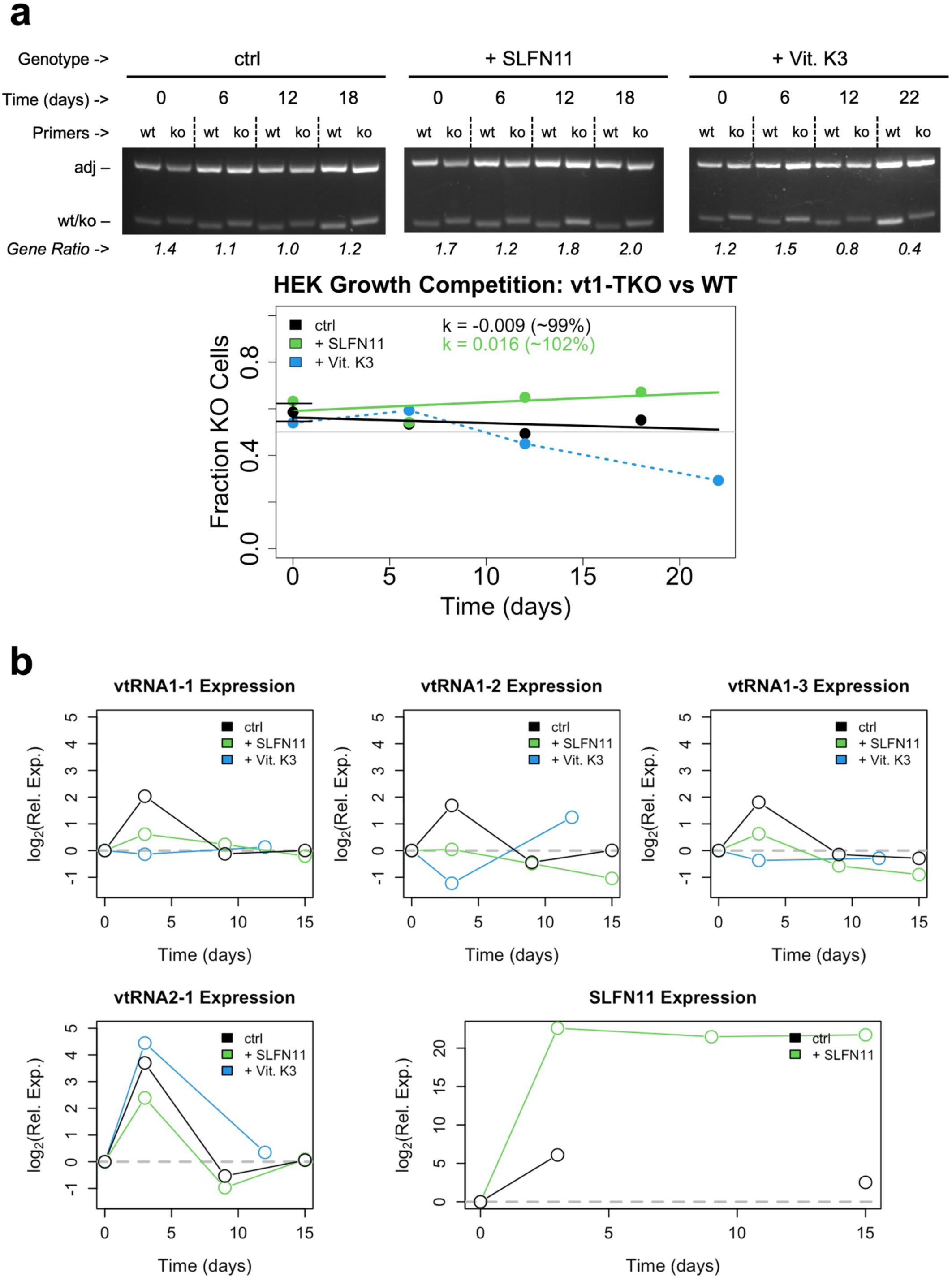
VTRNA1 knockout sensitizes HEK293T cells to menadione treatment but not SLFN11 overexpression. **[a]** Mixed populations of wild-type (WT) and vt1-TKO (c2) HEK293T cells were grown without treatment (“ctrl”, black), with repeated treatment of increasing menadione concentrations (“+ Vit. K3”, blue), or with SLFN11 overexpression (“+ SLFN11”, green). gDNA was sampled at the indicated time points, and the TKO:WT proportions tracked via PCR with indicated primer sets (“Primers”; see Figure 2a) and agarose gel electrophoresis. The prevalence of vt1-TKO versus WT cells (“Gene Ratio”) is the ratio of “ko” and “wt” product bands, each normalized to their respective “adj” product band. Plots of fraction KO cells versus time (bottom graph) for each triple-knockout clone were regressed (Eq. 1.1), and their rate constants (“k”, units of days^-1^) used to approximate relative growth rates for each clone (Eq. 1.2). Menadione data were not regressed (dashed line), since treatment was intermittent. **[b]** Quantification of vtRNA and SLFN11 relative RNA expression in mixed HEK293T experiments from panel-a. Values correspond to ΔΔC_q_, which is calculated as vtRNA/SLFN11 C_q_ normalized to GAPDH, then ΔC_q_ normalized to day-0 control (“ctrl”) for each RNA. Data points not connected by the trendline (e.g., SLFN11 expression in “ctrl”) indicate one or more intervening data points were expression was undetectable (C_q_ > 40). Panel-a and -b data are representative of two independent experiments.

SLFN11 overexpression did not disproportionately affect the survival/proliferation of *VTRNA1* triple-knockout cells (Figure 5a), nor did it alter vtRNA paralog expression in a consistent, meaningful manner (Figure 5b). During menadione (vitamin K_3_) treatment, while vtRNA paralog expression was similarly unaffected (Figure 5b), vt1-TKO cells appeared to have lowered survival relative to wild-type HEK293T cells (Figure 5a). However, we note that this sensitivity was only evident at higher menadione concentrations, where wild-type survival was also low. Based on the changes in cell line ratios as menadione concentration approached total lethality (Figure 5a), we estimate a ∼35% decrease in the vt1-TKO cells’ LC_50_ (menadione concentration at which there is 50% survival) relative to wild-type.

### RNA-sequencing of *VTRNA1*-knockout HEK293T cell lines

While vtRNAs did not appear to serve a critical function in HEK293T cells under standard, controlled growth conditions, the fact that they were expressed in wild-type cells suggested functionality. To probe for more subtle vtRNA-knockout effects, we isolated total RNA samples from our wild-type (HEK293T), CTRLdel, vt1-1 KO, vt1-2 KO, vt1-3 KO, and vt1-TKO cell lines and submitted them for (poly-A enrichment) RNA sequencing (Plasmidsaurus). The RNA samples from wild-type and single-knockout cell lines were also submitted for (rRNA depletion) RNA sequencing (Genewiz) for comparison. Differential expression analyses were performed relative to wild-type cells, since the CTRLdel cells – initially intended to serve as a control – turned out to have partially reduced vtRNA levels (Figure 3).

It’s prudent to note that the single- and triple-knockout/CTRLdel cell lines were from separate genetic editing experiments conducted months apart, making them susceptible to potential batch effects despite the shared protocol. Additionally, only ∼48% of the genes identified as differentially expressed (|log_2_FC| ≥ 1 (fold change) and FDR < 0.05 (false discovery rate)) from the single-knockout Plasmidsaurus sequencing data were also identified from the corresponding Genewiz data (Supplemental Figure 8a). This is not necessarily attributable to different sequencing methods, since RNA selection type and read depth also varied (see “RNA sequencing” methods). Finally, even within the Plasmidsaurus data, we observed high clonal variation in the differentially expressed genes (Supplemental Figure 8b). Hereafter, analyses/results are derived from the Plasmidsaurus data unless otherwise specificied.

There were broad transcriptomic changes in the vtRNA knockout cell lines, as well as the CTRLdel cells (Figure 6). A recent study (Stok et al. 2025) characterized the protein interactome of human vtRNAs by RNA-pulldown with mass spectrometry, offering us a more targeted approach for identifying relevant genes among those affected. Unfortunately, none of these “vtRNA binders” were differentially expressed in all three of our triple-knockout clones plus at least one of the single-knockout clones (Supplemental Figure 9) – our expected behavior for bonafide vtRNA-related genes. However, relaxing the criteria to account for batch effects between single- and triple-knockout cell lines, and for the CTRLdel cell line’s altered vtRNA expression, implicated genes might include H1-4, LGALS7, S100A16, and especially CRYAB. We note that genuine vtRNA-binding proteins would not necessarily have their expression affected by the loss of their RNA ligand, and thus, our relatively negative findings are not in conflict with the proteomic study (Stok et al. 2025).

**FIGURE 6.**
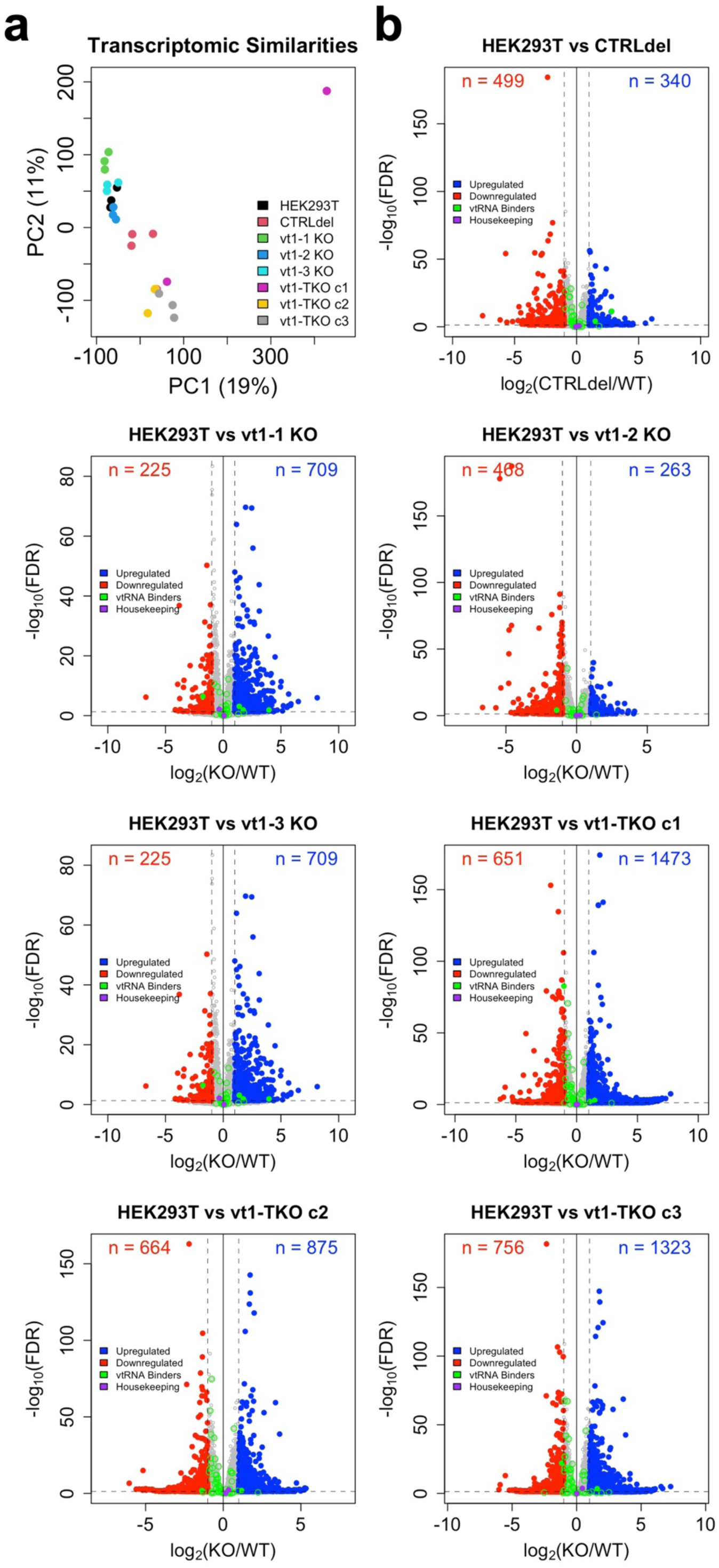
HEK293T cells lacking vtRNA1 paralogs have distinct transcriptomes. **[a]** Principal components analysis (PCA) of RNA-seq transcript counts per million transcripts (CPM) data for each HEK293T cell line. Data points correspond to RNA samples from independent biological replicates. **[b]** Volcano plots of transcripts differentially expressed between genetically edited cell lines and wild-type (WT) HEK293T. Upregulated (blue) and downregulated (red) transcripts have FDR < 0.05 (false discovery rate), and log_2_|FC| ≥ 1 (fold change), and their numbers are reported (“n”) for each differential expression analysis. The GAPDH and UBC transcripts (“Housekeeping”, purple) are shown as controls. Transcripts encoding proteins associated with vtRNA binding (“vtRNA Binders”, green) correspond to proteins previously identified by vtRNA pulldown (Stok et al. 2025).

Gene set enrichment analysis (GSEA) implicated numerous up- and downregulated biological processes across our various cell lines (Figure 7). Yet, none of these biological processes met our stringent criteria of being significantly affected in all three vt1-TKO cell lines, at least one single-knockout cell line, and not significantly affected in the CTRLdel cell line. By the more relaxed criteria described above, however, four annotated biological processes related to sensory perception are implicated.

**FIGURE 7.**
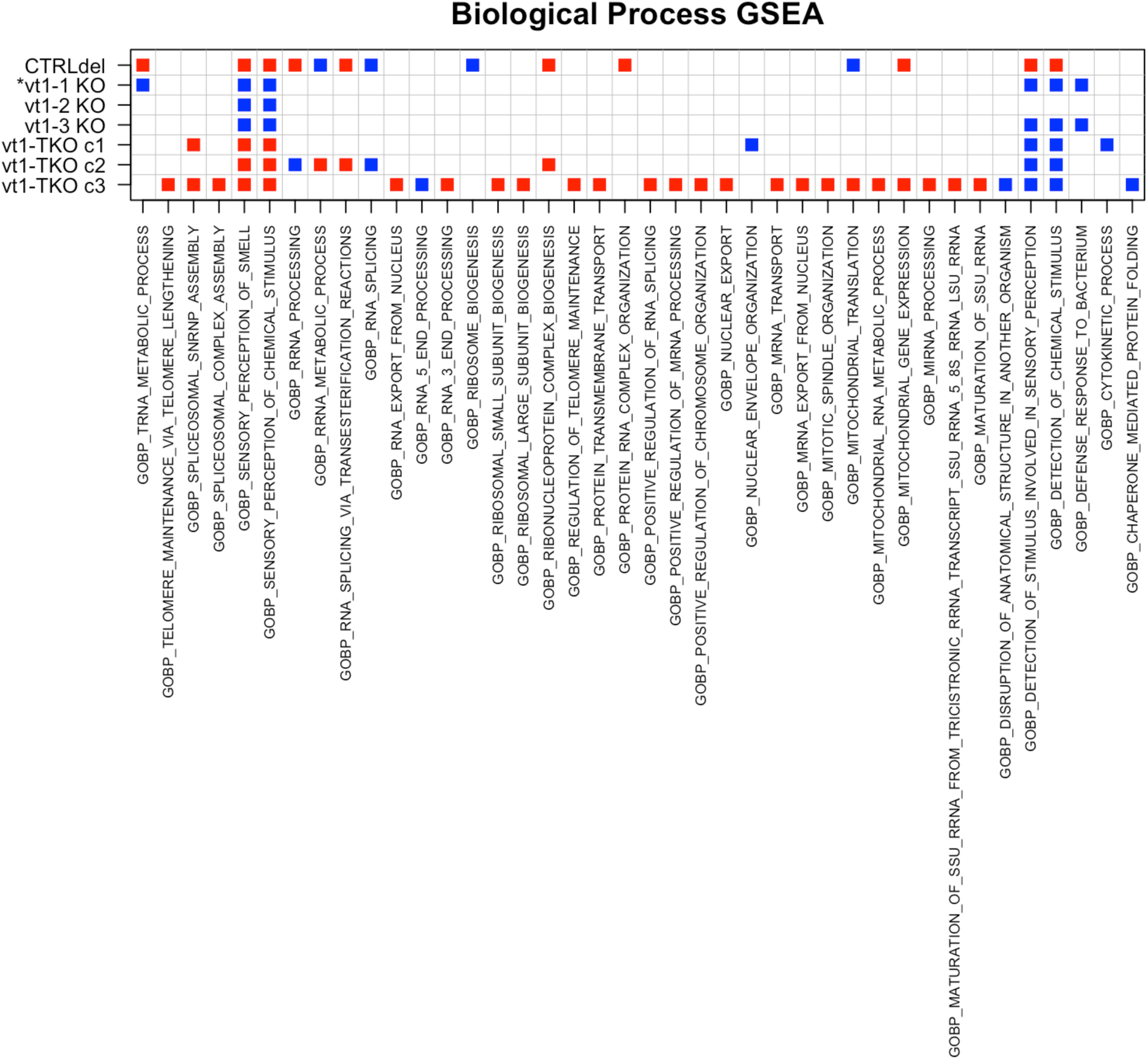
vtRNA1 paralogs knockout is correlated with varied biological processes in HEK293T cells. Gene Set Enrichment Analysis (GSEA) was performed using biological process gene ontology annotations (MSigDB set C5/GO, subset BP). Annotations significantly enriched among downregulated (red) or upregulated (blue) genes are shown for each differential expression data set. * The vt1-1 KO GSEA yielded an anomalous 321 statistically significant annotations, and the unique annotations were excluded for conciseness.

Focusing on individual genes whose expression is affected in all three vt1-TKO cell lines and at least one single-knockout cell line, there were 122 upregulated and 61 downregulated genes of interest. Gene ontology (GO) analysis implicated zero upregulated biological processes, and 16 downregulated biological processes related to filament/fiber organization and assembly, and to prostate gland development/morphogenesis. Further restricting the implicated genes to those not differentially expressed in the CTRLdel cell line, there were only 62 upregulated and 30 downregulated genes of interest. GO analysis of these genes implicated zero downregulated biological processes, and four upregulated biological processes related to nuclear localization of protein, binding, and smooth muscle cell differentiation. Perhaps in the future, a biological analysis will find a mechanistic connection between one of these processes and the vtRNAs.

## DISCUSSION

In the four decades since their discovery in association with vault particles, vtRNAs have been implicated in numerous biological processes. Yet, discrepancies between the findings of comparable studies, the seemingly paralog- and cell-specific nature of purported functions, and the lack of a phenotype in mice lacking vtRNAs have kept their molecular function(s) largely enigmatic (Prajapat et al. 2024; Taube et al. 2024; Aghajani Mir 2023; Gallo et al. 2022). Meanwhile, their functional relationship to their original namesake – vaults – has grown more ambiguous (Gallo et al. 2022; Nandy et al. 2009; Kickhoefer et al. 2002), while debate has arisen about the shared classification of human *VTRNA* loci and paralogs (Gallo et al. 2022; Lee et al. 2011; Stadler et al. 2009; Mosig and Stadler 2011; Im et al. 2020; Fort et al. 2018; Li et al. 2017; Lee 2015; Lee et al. 2014). Our findings shed light on the connection between the human vtRNA paralogs and on their cell-specific expression trends, but they also raise questions about vtRNA involvement in several previously implicated biological processes.

### Human *VTRNA* expression levels

Herein we report the numbers of molecules per cell of different vtRNAs for several human cell lines, using two orthogonal methods (Figure 1 and Table 1). Our findings indicate that *VTRNA1* paralog expression – both in terms of absolute and relative paralog levels – is reasonably consistent across tested cell lines, consistent with prior findings (van Zon et al. 2001; Kickhoefer et al. 1993). In contrast, we found that *VTRNA2-1* expression was much more variable, which appears attributable to *VTRNA2-1* epigenetic modifications, given that we found no gene sequence polymorphisms and that trans-acting factors necessary for vtRNA2-1 expression were present in HEK293T cells (Supplemental Figure 2-3). Based on prior studies, we suspect that the vtRNA2-1 expression differences among our cell lines may be due to differential DNA methylation within ∼200 bp upstream of *VTRNA2-1* (Raitoharju et al.; Yu et al. 2020; Romanelli et al. 2014; Rajić et al. 2025; Lee et al. 2014).

When they are expressed, human vtRNAs are observed at levels of a few thousand copies per cell (Table 1), which appears to be on the lower end for short ncRNAs (Table 2). Interestingly, this number is also dozens of times less than the generally cited range for the number of vault particles in human cells (Kickhoefer et al. 1998; Maniatis et al. 2025; González-Álamos et al. 2024; Frascotti et al. 2021). However, we note that the only (apparent) study quantifying vtRNA and vault particle levels in the same human cell line reported an excess of vtRNA molecules over vault particles (Kickhoefer et al. 1998), and we were unable to find a relevant, published quantification of vault particles per cell for any of our tested cell lines. The low copy number of the vtRNAs suggests that they function only under limited circumstances or with only a small subset of cellular proteins or nucleic acids, rather than having a housekeeping function in a general process like mRNA translation or pre-mRNA splicing (Table 2).

**TABLE 2.**
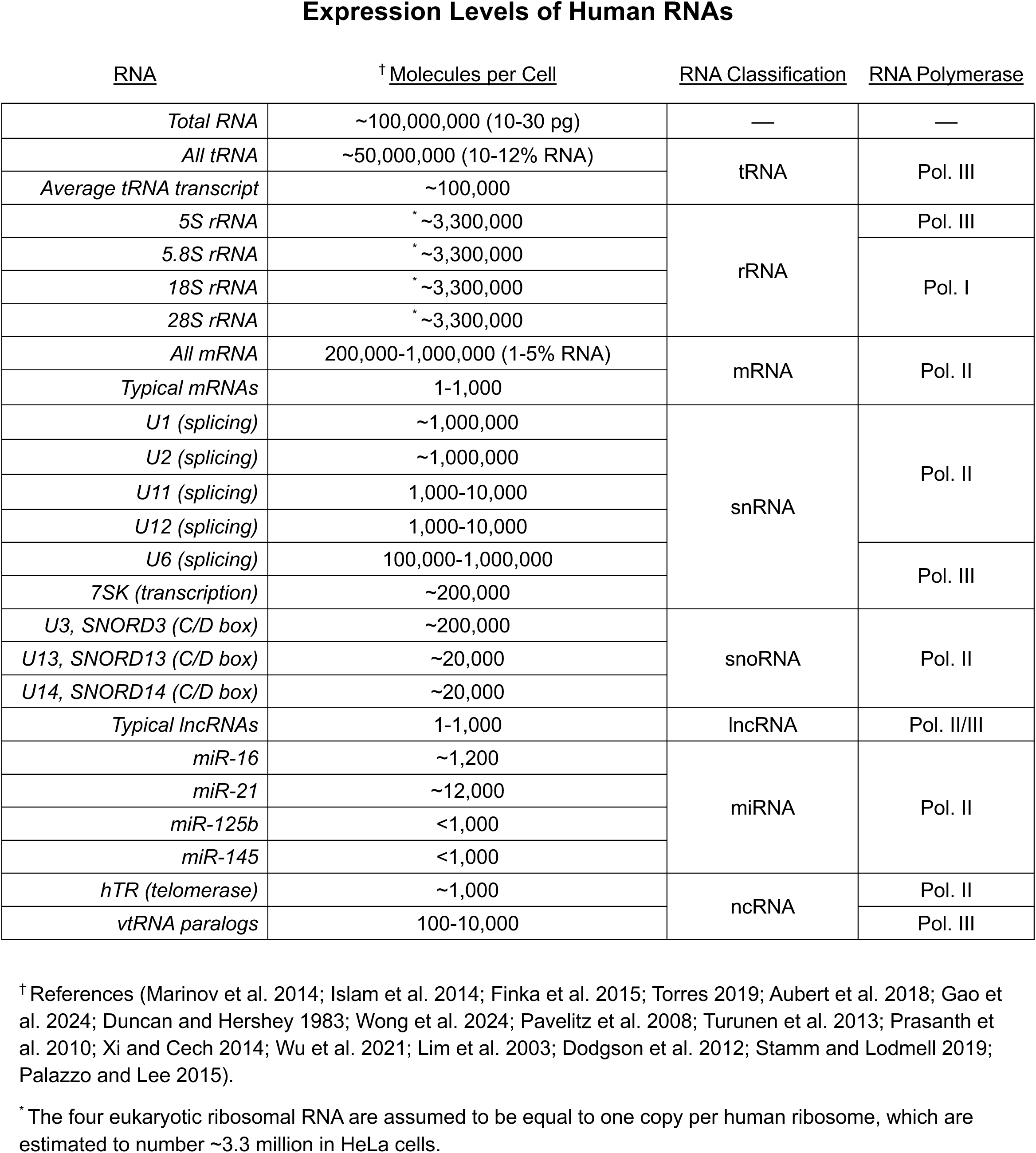
Expression levels of various human RNAs. Data are from the indicated references and refer to HEK293T or HeLa cell lines when possible. “Average” value for tRNA is the total number of all tRNAs per cell divided by the estimated number of different tRNA transcripts. “Typical” values of mRNAs and lncRNAs reflect the variation in copy number of specific transcripts of the indicated RNA class, sourced from associated references.

### Functional connection between human vtRNA paralogs

In humans, there are four functional vault RNA genes and two pseudogenes currently recognized: *VTRNA1-1*, *1-2*, and *1-3* (*VTRNA1* locus) and *VTRNA2-1* (*VTRNA2* locus) on chromosome 5q31, and *VTRNA2-2P* and *VTRNA3-1P* on chromosomes 2p14 and Xp11.2, respectively (van Zon et al. 2001; Stadler et al. 2009; Gallo et al. 2022). While *VTRNA1-2* and *1-3* are recent, lineage-specific duplications of *VTRNA1-1*, *VTRNA2-1* appears to be the result of a more ancient duplication event associated with the emergence of eutherians (placental mammals) (Stadler et al. 2009). While genetic evidence indicates that *VTRNA2-1* is a paralog of *VTRNA1*, its co-classification with the other human vtRNAs remains contentious, with many recent studies of vtRNA2-1 preferring to refer to it by its alternate name, nc886 (Rajić et al. 2025; Taube et al. 2024; Aghajani Mir 2023; Gallo et al. 2022; Im et al. 2020; Fort et al. 2020, 2018; Li et al. 2017; Lee 2015; Lee et al. 2014, 2011).

Key justifications for separate vtRNA2-1 classification include (1) *VTRNA2* lacking the TATA box and other external regulatory elements of *VTRNA1*, (2) much of *VTRNA2* transcript sequence similarity to *VTRNA1* being attributable to the internal Pol. III promoter box A/B motifs, (3) minimal association of vtRNA2-1 with the vault complex, (4) unique vtRNA2-1 expression patterns relative to *VTRNA1* paralogs, and (5) purportedly distinct *VTRNA2* and *VTRNA1* biological functions. However, it’s worth noting that association with the vault complex is no longer considered a defining feature even for the vtRNA1 paralogs, nor are the human vtRNA1 paralogs generally assumed to have a shared function (Gallo et al. 2022; Aghajani Mir 2023; Taube et al. 2024). Thus, any functional connection between human vtRNAs, including vtRNA2-1, remains an open question.

Our expression studies in HEK293T wild-type and vtRNA-knockout cells (Figure 3) revealed no credible evidence that vtRNA knockout leads to compensatory expression of remaining *VTRNA1* paralogs, consistent with the idea that *VTRNA1* paralogs are not purely redundant. However, significantly increased vtRNA2-1 expression was a very consistent observation across our *VTRNA* single- and triple-knockout cell lines, suggesting that *VTRNA1* and *VTRNA2* paralogs may not perform wholly independent functions *in vivo*, even if their molecular mechanisms are distinct. Notably, a recent study utilizing *VTRNA1* triple-knockout A549 cells observed similar inter-locus vtRNA compensation, though the effects were less exaggerated and went unacknowledged by the study’s authors (Stok et al. 2025).

Of course, we acknowledge that such an expression correlation does not necessarily equate to a cross-paralog regulatory mechanism and/or shared function between the vtRNA paralogs. As examples, similar inter-locus correlation could be produced if the modified *VTRNA1* sequence overlapped with an enhancer for *VTRNA2-1*, if the loss of transcription at an edited gene perturbed insulation effects around the differentially expressed one (Brasset and Vaury 2005; Gerasimova and Corces 2001), or if the physical process of genetic editing/selection itself had an indirect effect on *VTRNA2-1*. However, there are no annotated *VTRNA2-1* enhancers affected by our *VTRNA1* deletions, the *VTRNA1* and *VTRNA2* loci are too distant for typical insulation effects, and the CTRLdel cells do not exhibit vtRNA2-1 upregulation. Thus, given the consistent upregulation we observed and the shared evolutionary history of the vtRNA1 and 2 paralogs, a regulatory mechanism connecting their functions/expression seems plausible and warrants further investigation.

The natural next step would of course be interrogating the phenotype of a quadruple knockout cell line that also lacks vtRNA2-1. However, we note that vtRNA2-1 is not detectably expressed in wild-type HEK293T cells (Figure 1), and that despite its apparent upregulation in our vtRNA1 knockout cell lines (Figure 3), vtRNA2-1 absolute expression levels are still quite low – less than 100 cpc. Thus, pursuing such a quadruple knockout in HEK293T cells seems unlikely to yield a compelling result. Fortunately, our findings indicate that both HeLa and MRT cells have robust vtRNA2-1 expression with identical *VTRNA* gene sequences to HEK293T cells (Figure 1 and Supplemental Figure 2). Perhaps future research will exploit this uniquely tractable opportunity to apply our efficient *VTRNA1* knockout strategy to generate vtRNA quadruple-knockout MRT cells and interrogate their phenotype with our highly sensitive growth competition assay.

### Biological role of vtRNAs

Multiple studies have implicated *VTRNA1* paralogs in contributing to cell proliferation, especially in cancer cells, and reported *VTRNA1* single-knockout human cell lines with growth defects (Gallo et al. 2022). A recent study even generated *VTRNA1* single-knockouts in HEK293T cells, much like this study, and reported a growth deficit (Alagia et al. 2023). Yet, our findings clearly suggest no significant, basal growth deficit in HEK293T cells lacking *VTRNA1* paralog expression (Figure 4a). We note that, while most comparable studies monitored absolute growth with isolated, homogenous cell populations, we favored a competition-based assay with heterogeneous cell populations to quantify relative growth rates while controlling for variability in growth conditions. As a result, it’s possible to reconcile these disparate findings if, for example, the wild-type cells compensated for a hypothetical cytokine deficiency in vtRNA-knockout cells in our co-culture assays.

Our experiments interrogating the effects of *VTRNA1* knockout on menadione (vitamin K_3_) sensitivity (Figure 5) suggested that vtRNAs can protect against oxidation-induced cell death, consistent with their potential implication in RNA repair (Burroughs and Aravind 2016; Akiyama et al. 2022). However, it’s prudent to emphasize that the effect was quite modest (∼35% lower LC_50_), calling biological relevance into question. On the other hand, we note that a similar magnitude of menadione sensitivity was reported for RtcB (Akiyama et al. 2022), which is known to function in tRNA maturation/repair, presumably because menadione’s broad effects make it difficult for a single gene to offer more significant protection (Loor et al. 2010; Morrison et al. 1985). Regardless, *VTRNA1* knockout cells were not sensitized to SLFN11 overexpression (Figure 5), suggesting that any menadione sensitivity is not attributable to a vtRNA role in that RNA repair pathway.

Interestingly, these effects might be explained if vtRNAs function as negative regulators of apoptosis, as implied in prior studies (Gallo et al. 2022; Taube et al. 2024; Aghajani Mir 2023; Büscher et al. 2022; Bracher et al. 2020; Horos et al. 2019). In such a case, vt1-TKO and wild-type cells could be equally susceptible to the molecular damage of potent oxidation, but cells lacking vtRNAs trigger apoptosis more readily past that damage threshold, resulting in a modest yet detectable survival bias at moderately lethal menadione concentrations. Unfortunately, our flow cytometry data does not indicate increased apoptosis in *VTRNA1* knockout cells under normal growth conditions (Figure 4b and Supplemental Figure 5).

vtRNAs have also been implicated in innate immunity, specifically during infection of various human cell lines by several RNA viruses (Stok et al. 2025; Aghajani Mir 2023; Gallo et al. 2022; Nandy et al. 2009; Peng et al. 2016). Yet, we found no evidence that vtRNA1 paralogs meaningfully impact viral infection or cell viability during OC43 coronavirus infection of HEK293T cells (Supplemental Figure 6). Naturally, we cannot discount cell- and/or virus-specific effects.

Finally, our RNA sequencing data (Figures 6-7 and Supplemental Figure 8-9) did not convincingly implicate vtRNAs in any biological processes, nor did they seem to recapitulate existing transcriptomic analyses of vtRNA-knockout cell lines (Gallo et al. 2025; Alagia et al. 2023). However, our studies differed in multiple meaningful ways – RNA enrichment and sequencing methods, vtRNA knockout strategies, cell types, alignment and differential expression software, ontology analyses, etc. – which could account for our incongruent findings.

Overall, our data indicate that vtRNA1 paralogs do not serve a unique, critical function in HEK293T cells under standard growth conditions. Furthermore, while the human *VTRNA1* paralogs have been previously implicated in biological processes such as proliferation/growth, apoptosis/autophagy, viral infection, and more (Taube et al. 2024; Aghajani Mir 2023; Gallo et al. 2022), several of our key findings are inconsistent with vtRNAs playing a major role in many of these processes.

### Concluding remarks

Despite decades of focused study, the biological role(s) of human vtRNAs and their functional relationships (or lack thereof) to one another remain enigmatic. Our findings indicate that, while the human *VTRNA1* versus *VTRNA2* paralogs indeed exhibit distinct expression patterns and may have distinct functions/mechanisms, there is also a potential regulatory mechanism connecting their expression, suggesting that the vtRNA paralogs’ biological roles are not completely independent/unrelated. In addition, our findings call into question the generalizability of certain purported human vtRNA functions, reinforcing the need for further, critical study of vtRNA biological function(s).

## MATERIALS & METHODS

### CRISPR-Cas9 knockout of *VTRNA* genes

Each gene deletion required two plasmids, each expressing Cas9 and one of the requisite sgRNAs. The pX459 vector (Addgene #62988) for combined Cas9 and sgRNA expression was cleaved with BbsI restriction enzyme (NEB #R3539) according to manufacturer specifications. Respective complementary oligos (Supplemental Table 1, “sgRNA”) were annealed in annealing buffer (50 mM TRIS pH 7.5, 200 mM NaCl) with a 3 h, 95➔20°C thermocycler gradient. BbsI-cut pX459 was ligated to the annealed inserts with T4 DNA ligase (NEB #M0202) according to manufacturer’s instructions.

HEK293T cells were grown (see “culture of human cell lines”) in a T25 (Corning #430639) to ∼80% confluency. Cells were washed twice with media lacking antibiotics, then left in 4 mL media without antibiotics. In parallel, 600 ng each of site “a” and “b” pX459 plasmids (see above, Figure 2, Supplemental Table 1) were suspended in 500 µL Opti-MEM (Gibco), while 20 µL of Lipofectamine 2000 (Invitrogen #11668027) was suspended in 500 µL Opti-MEM and incubated at room temperature for 5 min. These two transfection solutions were combined and incubated for 20 min at room temperature, then added to the cell media. Cells were left to transfect for 6 h, then changed to standard media and cultured overnight.

At ∼24 h post-transfection, 4 µg/mL puromycin was added to cell media. Puromycin selection was continued for 3 d, with daily media/puromycin exchange. After, cells were trypsinized and passaged into a fresh T25, then grown under standard conditions to 80-90% confluency. Next, cells were trypsinized and suspended in PBS (phosphate buffered saline) with 2% FBS (fetal bovine serum) at 10^6^ cells/mL, then sorted with a FACSAria Fusion Cell Sorter (BD Biosciences) with 100 µm nozzle. FSC and SSC were used to sort singlets into 96-well plates (Corning #3596) with 200 µL conditioned media, then clones were cultured for ∼14 d. Clones were sampled and initially genotyped via PCR with Q5 Blood Direct 2X Master Mix (NEB #M0202) and agarose gel electrophoresis.

For validated clones in Figure 2, genomic DNA (gDNA) was isolated with the DNeasy Blood & Tissue Kit (Qiagen #69504), PCR amplified with Q5 High-Fidelity 2X Master Mix (NEB #M0492) and the indicated primer sets (Figure 2a and Supplemental Table 1), and visualized by ethidium bromide staining of 2.5% agarose gels, according to respective manufacturer protocols.

### Culture of human cell lines

Unless otherwise specified, HEK293T (ATCC #CRL-11268, Lot #63696280), HeLa (ATCC #CCL-2), and U2OS (ATCC #HTB-96, Lot #70058619) cells were incubated at 37°C, 5% CO_2_, and ≥80% humidity in DMEM (Gibco) with 10% v/v FBS, 1X GlutaMAX (Gibco), and 1% w/v penicillin/streptomycin. Human BJ skin fibroblasts (ATCC #CRL-2522, Lot #70063200) were incubated at 37°C, 5% CO_2_, and ≥80% humidity in EMEM (ATCC) supplemented with 10% FBS, 1X GlutaMAX (Gibco), and 1% w/v penicillin/streptomycin. Human MRT / G-401 kidney tumor cells (ATCC #CRL-1441, Lot #70055408) were incubated at 37°C, 5% CO_2_, and ≥80% humidity in McCoys 5A (ATCC) supplemented with 10% FBS, 1X GlutaMAX (Gibco), and 1% w/v penicillin/streptomycin. Human induced pluripotent stem cells (iPSCs) were a gift from the Roy Parker or John Rinn labs (CU Boulder).

### In vitro transcription (IVT)

1 mL reactions were prepared in RNA Polymerase buffer (40 mM TRIS pH 8.0, 6 mM MgCl_2_, 1 mM DTT, 2 mM spermidine) supplemented with 1.5 mM each rNTP mix (Promega #E6000), an additional 22 mM MgCl_2_, 90 mM DTT, 0.1% v/v RNase inhibitor (Promega #N2618), 2 µM Milligan template DNA (“template” in Supplemental Table 1) (Milligan and Uhlenbeck 1989), and ∼140 nM T7 RNA polymerase, then incubated overnight at 37°C.

Completed reactions were centrifuged 16,000*g* for 10 min at room temperature to remove pyrophosphate, then supernatant collected and treated with 0.5% v/v Turbo DNase (Invitrogen AM2238) at 37°C for 15 min. Next, equal volume phenol:chloroform was added, samples vortexed thoroughly and centrifuged 16,000*g* at room temperature for 3 min, and the aqueous phase collected, then these steps repeated with chloroform:isopentanol. Phenol-chloroform-extracted RNA was precipitated by sequential addition of 300 mM sodium acetate and 75% ethanol, followed by 1 h incubation at -80°C and several repeated 75% ethanol wash cycles via centrifugation (16,000*g* at 4°C for 30 min) and supernatant removal.

RNA pellets were resuspended in sequencing loading dye (formamide with 20 mM EDTA, 0.1% v/v bromophenol blue, 0.1% v/v xylene cyanol) and resolved on a (8 M, 7%) urea-polyacrylamide gel (1 h electrophoresis, constant 50 mA, 1X Tris borate EDTA (TBE) buffer), then full-length product bands excised under UV shadowing. Gel fragments were incubated overnight at room temperature in 10 mM TRIS pH 7.0, then supernatant subjected to ethanol precipitation as described above. RNA pellets were resuspended in 10 mM TRIS pH 7.0 and stored at -80°C. RNA concentrations were determined via A_260_ by NanoDrop (Thermo Fisher).

### Cellular RNA purification

HEK293T, HeLa, U2OS, MRT, and BJ fibroblast cells were cultured under their respective standard conditions in T25 flasks (Corning #430639) until ∼80% confluency, then washed with PBS, lysed in 1 mL TRIzol (Invitrogen), and their RNA purified (with DNase treatment) via the Direct-zol RNA Miniprep Plus kit (Zymo Research #R2070) according to manufacturer guidelines. For human iPSCs, RNA samples were a gift from the John Rinn lab (CU Boulder), or frozen cell pellets were a gift from the Roy Parker lab (CU Boulder) then lysed with TRIzol and subjected to the same RNA purification protocol as cultured cells.

### RT-qPCR

cDNA was prepared from purified RNA with the LunaScript RT SuperMix kit (NEB #E3010), and qPCR was performed with Luna Universal qPCR Master Mix (NEB #M3003) and the indicated primers (“probes”, Supplemental Table 1), both as directed. A melt curve was generated following amplification using a 60➔95°C gradient at 2 °C/min.

For copies per cell (cpc) calculations, concentration standards of IVT products were prepared by serial dilution, then used in cDNA reactions in known quantities relative to cellular RNA reactions. Plots of quantification cycle (C_q_) versus standard concentrations were fit with linear regression, then used to interpolate the relative mass of vtRNA within cellular RNA. Mass proportions were converted to cpc via multiplication by Avogadro’s number (6.022x10^23^ mole^-1^) and RNA mass per cell (20 pg) (Russo et al. 2014), and division by the respective vtRNA molecular weight (vtRNA1-1 = 31,767 Da, vtRNA1-2 = 28,525 Da, vtRNA1-3 = 28,401 Da, vtRNA2-1 = 31,960 Da).

For relative expression calculations, vtRNA (or SLFN11) C_q_ values were first normalized to GAPDH C_q_ values within each sample, then ΔC_q_ values were normalized to the average ΔC_q_ value of the wild-type (or day-0 control) samples to give log_2_ fold changes.

### Northern blots

Probes (Supplemental Table 1, “probes” bottom sequences) and pBR322 MspI ladder (NEB #N3032) were dephosphorylated, then radio-labeled in reactions containing 1X kinase buffer (NEB #B0201), 0.5 U/µL PNK enzyme (NEB #M0201), 20 µM oligo or 5% v/v ladder, and 50 µM γ-^32^P-ATP. Labeled DNA was purified with Micro Bio-Spin P-6 columns (Bio-Rad #7326221) according to product instructions.

1.5 mm thick, 28x20.5 cm urea-polyacrylamide (7 M / 6%) gels were prepared in 1X TBE. RNA samples and ladder were diluted in 2X loading dye (93% v/v formamide, 30 mM EDTA, 0.1X TBE, 0.05% bromophenol blue and xylene cyanol) and heated at 95°C for 5 min. Following pre-running the gel at constant 40 W for 30 min, samples were loaded and run at constant 40 W for 2 h. Gels were trimmed and placed into a 0.5X TBE-soaked transfer sandwich with Whatman 3MM CHR paper (Cytiva #3030-917) and Hybond-N+ membrane (Cytiva #RPN203B), then electrophoresed at 4°C and constant 1.5 A for 1.5 h in 0.5X TBE.

Membranes were crosslinked on a CL-1000 ultraviolet crosslinker (Krackeler Scientific) at 254 nm with 1200x100 µJ/cm^2^. Crosslinked membranes were wet with PerfectHyb PLUS (Sigma-Aldrich #H7033) hybridization buffer, then placed into glass hybridization tubes with 25 mL hybridization buffer and incubated in a rotator at 42°C for 30 min. 1x10^7^ cpm of radio-labeled probes were suspended in 1 mL hybridization buffer and heated at 95°C for 5 min, added to the hybridization tube, then incubated overnight at 42°C with rotation. After, membranes were rinsed with 75 mL 2X SSC (1X = 150 mM NaCl, 15 mM sodium citrate, pH 7.0) + 0.1% w/v SDS and washed three times with 75 mL 2X SSC + 0.1% SDS via 15 min incubation at 42°C with rotation, rinsed and washed once with 0.2X SSC + 0.1% SDS, then rinsed and washed once with 0.1X SSC + 0.1% SDS.

Blots were dried and exposed on phosphor screens for 1 h to 12 d, then screens imaged on a Typhoon FLA 9500 (GE). Densitometry was performed with ImageQuant TL v10.2 (Cytiva) and counts per cell (cpc) calculations were performed similarly to the method described in the “RT-qPCR” section.

### Competitive growth assays

6x10^5^ cells each of wild-type and a vt1-TKO clone were seeded into T25 flasks (Corning #430639), then ∼85% of cells immediately taken as day-0 samples. After, mixed cultures were grown under standard conditions for the indicated lengths of time with regular passaging every ∼3 d. At each passage, excess cells (∼85%) were taken as culture samples. Samples were gDNA purified with the DNeasy Blood & Tissue Kit (Qiagen #69504), PCR amplified with Q5 High-Fidelity 2X Master Mix (NEB #M0492) and the indicated primer sets (Figure 2a and Supplemental Table 1), and visualized by ethidium bromide staining on 2.5% agarose gels, according to respective manufacturer protocols.

Densitometry was performed on agarose gels with GelAnalyzer v19.1, then “wt” and “ko” primer-associated band intensities normalized to their respective “adj” primer-associated band intensities (see Figure 2 for notation). Normalized “ko” band intensities were divided by the sum of “wt” and “ko” normalized band intensities to give the metric of “Fraction KO Cells” (Figure 4). Plots of this metric versus time were regressed with Eq. 1.1,

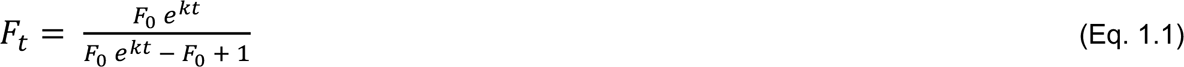

where *F_t_* is the knockout cell fraction at a given time, *F_0_* is the initial knockout cell fraction, *t* is time, and *k* is the difference in the vt1-TKO clone’s and wild-type’s growth rate constants (*k* = *k_ko_* - *k_wt_*). Relative growth rates were estimated from the fitted *k* values with Eq. 1.2,

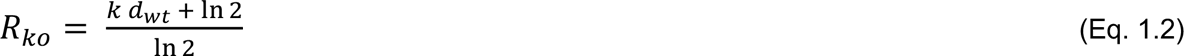

where *R_ko_* is the vt1-TKO clone’s growth rate relative to wild-type, *k* is the fitted rate constant value from Eq. 1.1 in units of d^-1^, and *d_wt_* is the doubling time for wild-type cells, assumed to be 0.75 d for these calculations.

The p-value and 95% confidence interval in Figure 4a are from a one-sample t-test (mu = 0) of the regressed Eq. 1.1 *k* rate constants from each vt1-TKO clone, using the “t.test” function in base R.

### RNA sequencing

Three RNA samples (biological replicates) for each indicated cell line were submitted to Plasmidsaurus (Louisville, Kentucky USA) and Genewiz (South Plainfield, New Jersey USA) for RNA sequencing. The former utilized poly(A) enrichment and 3’-end Illumina sequencing with ∼10 million single-end reads, and the latter utilized rRNA depletion and Illumina sequencing with ∼50 million paired-end reads. For the Plasmidsaurus sequencing, each RNA sample (biological replicate) was submitted in triplicate (technical replicates) and FASTQ data pooled to artificially triple the single-end read depth to ∼30 million. RNA-seq data were aligned and analyzed in R v4.5.2 with custom scripts that have been uploaded to GitHub (github.com/whemphil/vtRNA_Manuscript), and raw sequencing data have been uploaded to the Gene Expression Omnibus (GSE335700).

Briefly, index building was performed with the “BSgenome.Hsapiens.UCSC.hg38” package, FASTQ quality filtering with the “Rfastp” package, alignment and feature counting with the “Rsubread” package “subjunc” and “featureCounts” functions and Ensembl release 115 GRCh38 GTF file, and differential expression analysis with the “DESeq2” package. Gene ontology and gene set enrichment analyses were performed using the MSigDB C5-BP annotations via the “msigdbr” package. Proteins implicated as part of the vtRNA interactome were taken from RNA-pulldown mass spectrometry data in (Stok et al. 2025).

### Gene fragment sequencing and alignment

Human iPSC cell pellets were generously provided by the Roy Parker lab (CU Boulder), and other cell lines were cultured as described (see “culture of human cell lines”), then their gDNA was isolated with the DNeasy Blood & Tissue Kit (Qiagen #69504) according to the manufacturer protocol. vtRNA gene fragments were amplified from each gDNA sample with Q5 High-Fidelity 2X Master Mix (NEB #M0492) according to product recommendations. For wild-type and single-knockout cell lines, separate PCR reactions were performed with the vt11, vt12, vt13, and vt21 primer sets, and for the triple-knockout cell lines, the ko and vt21 primer sets were used (Figure 2a and Supplemental Table 1).

Samples of completed PCR reactions were submitted to Plasmidsaurus (Louisville, Kentucky USA) for sequencing, and the results were aligned with SnapGene v4.2.11 and the Clustal Omega Multiple Sequence Alignment Tool (EMBL-EBI).

### Flow cytometry

1x10^5^ cells of the indicated HEK293T lines were seeded into T25 flasks (Corning #430639) and cultured under standard conditions for 3 d. Culture supernatant and trypsinized cells were pelleted together and resuspended to 10^6^ cells/mL in PBS, then 100 µL samples were stained with the Annexin V Apoptosis Kit (Novus Biologicals #NBP2-29373) according to manufacturer instructions.

Stained samples were analyzed on a BD Accuri C6 PLus Flow Cytometer analyzer (BD Biosciences), and the Floreada web server (https://floreada.io) was used for data processing and gating. Gating strategy was (1) SSC-A/FSC-A to remove debris, (2) FSC-A/FSC-H to identify singlets, then (3) PerCP-A/FITC-A (i.e., PI/Annexin-V) to partition cells into live, apoptotic, and dead fractions (Supplemental Figure 5). Compensation controls for PI/Annexin-V gate were wild-type cells stained with PI only or Annexin-V only.

### OC43 viral infection assays

10^5^ cells were seeded into each well of a 6-well plate (Corning #3516) and incubated under standard conditions for 24 h. For viral replication experiments, cells were homogeneous populations of wild-type or vt1-TKO clones, and for viability/growth experiments, cells were an equal mix of wild-type and a vt1-TKO clone. At ∼24 h post-seeding, cells were washed once with DMEM (Gibco) containing 2% v/v FBS (infection media) and inoculated with OC43 at MOI 0.05 in 1 mL of infection media for 1 h. After viral absorption, cells were washed once with infection media and placed into 2.5 mL of infection media for the duration indicated.

For viability/growth experiments, cells were trypsinized and pelleted at 1, 2, or 3 d post-infection, then the proportion of wild-type versus knockout cells determined as described (see “competitive growth assays”).

For viral replication assays, at 1 d post-infection cells were washed with PBS and lysed in TRIzol (Invitrogen), then RNA purified with the Direct-zol RNA Miniprep Plus kit (Zymo Research #R2070) according to manufacturer guidelines. cDNA was prepared using the High-Capacity cDNA Kit (ThermoFisher #4368814) with 1 µg of input RNA, according to the manufacturer’s instructions. qPCR was performed with iQ SYBR Green (Bio-Rad #1708880) in a 10 µL reaction with 125 nM of each qPCR primer (Supplemental Table 1) and 1.25 ng equivalents of cDNA. Thermocycler conditions were (1) 95°C for 2 min, followed by (2) 40 cycles of 95°C for 15 s then 60°C for 30 s. A melt curve was generated following amplification using a 55➔95°C gradient at 0.5 °C/s, which revealed no off-target amplification.

For relative expression calculations, OC43 gRNA C_T_ values were first normalized to GAPDH C_T_ values within each sample, then ΔC_T_ values were normalized to the average ΔC_T_ value of the wild-type samples to give ΔΔC_T_ values. Reported relative expression values are 2^(ΔΔC_T_).

### Ectopic expression of vtRNAs

Complementary DNA oligos (Supplemental Table 1, “insert”) coding for vtRNA1-1 and vtRNA2-1 – with 40 bp of flanking sequence – were annealed and cloned into the HinDIII-EcoRI site of the polylinker region of pUC19. Complementary DNA oligos (Supplemental Table 1, “U6 insert”) coding for vtRNA1-1 were annealed and cloned into the BbsI site of pJR255 (Addgene #78549). Plasmids were transfected into HEK293T cells without any chemical selection (see “CRISPR-Cas9 knockout of VTRNA genes”), and RNA isolated at 48 h post-transfection (see “Cellular RNA purification”).

### SLFN11 overexpression and menadione treatment assays

6x10^5^ cells each of wild-type and a vt1-TKO clone were seeded into T25 flasks (Corning #430639), then ∼85% of cells immediately taken as day-0 samples. After, mixed cultures were grown under standard conditions with repeated treatment/transfection and cell sampling. On treatment/transfection days, cells were washed with Opti-MEM (Gibco), treated/transfected in Opti-MEM for 4 h (just Opti-MEM for controls, “ctrl”), then low-serum media replaced with standard media. On sample days, cells were trypsinized, ∼85% of cells aliquoted and pelleted, and remaining cells re-seeded in standard media.

For SLFN11 overexpression experiments, 1 µg of SLFN11-expressing plasmid (OriGene #RC226247) was transfected into cells without any chemical selection (see “CRISPR-Cas9 knockout of VTRNA genes”) on days 2, 5, 8, 11, 14, and 17. Samples were taken on days 3, 6, 9, 12, 15, and 18.

For menadione (vitamin K_3_) experiments, cells were treated with 10, 20, 25, or 30 µM menadione (Sigma-Aldrich #47775) on days 2, 5, 8, and 14, respectively. Samples were taken on days 3, 6, 12, and 22. Treatment-to-sampling intervals exceeding 1 d were due to excessive cell death at high menadione concentrations, which required extended cell recovery time for adequate samples.

For survival/proliferation assays (Figure 5a), cell samples were used for gDNA isolation, PCR amplification with the indicated primers (“wt”, “ko”, and “adj”; Supplemental Table 1), and gene ratio visualization via agarose gel electrophoresis (see “Competitive growth assays”). For RNA expression (Figure 5b), cell samples were resuspended in 350 µL TRIzol (Invitrogen), total RNA isolated (see “Cellular RNA purification”), and RT-qPCR performed (see “RT-qPCR”) with indicated primers (“probes”, Supplemental Table 1). For SLFN11 protein expression (Supplemental Figure 7c), cell samples were resuspended in 1X sample buffer (Novex #NP0008) and used for western blots (see “Western blots”).

For experiments to validate SLFN11 overexpression (Supplemental Figure 7a-b), 4x10^5^ wild-type cells were seeded in T25 flasks, transfected the next day as described above (Opti-MEM only for controls, “ctrl”), then trypsinized and sampled at 1 and 2 d post-transfection. For protein expression (Supplemental Figure 7a), cell samples were lysed in 1X RIPA buffer (20 mM Tris pH 7.5, 150 mM NaCl, 1% v/v NP-40, 1% w/v sodium deoxycholate, 0.1% w/v SDS, 1 mM EDTA) plus Complete Protease Inhibitor (Thermo Scientific #A32965), mixed with an equal volume of 2X sample buffer (Novex #NP0008), then used for western blots (see “Western blots”). For RNA expression (Supplemental Figure 7b), cell samples were resuspended in 350 µL TRIzol (Invitrogen), total RNA isolated (see “Cellular RNA purification”), and RT-qPCR performed (see “RT-qPCR”) with indicated primers (“SLFN11 probes”, Supplemental Table 1).

### Western blots

Samples were run (200 V, 40 min) on 4-12% acrylamide Bis-Tris gels (Invitrogen #NP0321BOX) in 1X MOPS SDS Running Buffer (Invitrogen #NP0001) with a PageRuler Plus Prestained Protein Ladder (ThermoScientific #26619). Gels were transferred (0.5 A, 1 h, 4°C) to Hybond ECL membrane (GE #RPN78D) in 1X western buffer (25 mM Tris pH 7.5, 192 mM glycine, 0.1% w/v SDS, 20% v/v methanol).

Membranes were incubated for 15 min (25°C, 60 rpm) in blocking buffer (ThermoScientific # 37539), then incubated overnight (4°C, 60 rpm) with a 1:1000 dilution (in blocking buffer) each of rabbit α-SLFN11 (Abcam #ab121731) and/or rabbit α-GAPDH (Cell Signaling Technology #2118S) primary antibodies.

Membranes were washed 3x with PBS + 0.05% v/v Tween 20 and 1x with PBS (15 min, 25°C, 60 rpm), then incubated 1.5 h (25°C, 60 rpm) with a 1:1000 dilution (in blocking buffer) of goat anti-rabbit IgG-HRP secondary antibody (Santa Cruz Biotechnology #Sc2301). Membranes were washed again, incubated 5 min (25°C, 60 rpm) in SuperSignal West Pico PLUS solutions (ThermoScientific #34580), then imaged on a FluorChem R system (Bio-Techne) with a 5 s to 2 min exposure.

### Diagram, reaction scheme, and figure generation

Tables were prepared with Word (Microsoft), graphs were prepared with R v4.5.2, and figures and schematics were assembled in PowerPoint (Microsoft).

## ACKNOWLEDGEMENTS

W.O.H. was supported by the National Institutes of Health (F32 GM147934). T.R.C. is an investigator of the Howard Hughes Medical Institute. We thank the John Rinn lab (University of Colorado Boulder) for providing the human iPSC RNA and for stimulating discussion and feedback throughout these studies, as well as Mary Allen and Robin Dowell (University of Colorado Boulder) for their helpful insights throughout. We also thank Theresa Nahreini, Emily Proksch, the Flow Cytometry Shared Facility in JSCBB (RRID:SCR_019309; NIH grant S10OD021601), and the Biochemistry Cell Culture Facility (RRID:SCR_018988) for technical assistance and equipment use during these studies.

## COMPETING INTERESTS STATEMENT

T.R.C. is a scientific advisor for Eikon Therapeutics. The other authors have no competing interests to declare.

## Notes

### Summary of Updates

As an alternative to generating more knockout cell lines, we have added an additional paragraph at the end of the 'Functional connection between human vtRNA paralogs' Discussion section to address the reason for us not examining a quadruple vtRNA knockout.

