## Supplementals for "Human vault RNAs exhibit diverse expression patterns and inter-locus compensation"

### Synthetic Oligonucleotides Used

| *Name* | *Sequence (5' -> 3')* |
|---|---|
| vt11 | ACTCTCTTTTTGACTCACTCTTTGTGACG |
| | TCCTTTTACCATGGAAAGGGAGCTG |
| vt12 | ACTGTTGTTTTGACTCACTTTGTGACG |
| | CTCTGTATTTCCTGCCCAGTGCTAAAATC |
| vt13 | CACCGTTGTTTTGACTCACTCTTTGTG |
| | CTGCCCACTACTAAAATCAGCCATTTC |
| vt21 | GGAGTTAGCTCAAGCGGTTACCT |
| | CCCAGCACAGAGATGGACAGATAG |
| wt | AAGGAATCAGGGCTCCTCGG |
| | GGGCCTCAGTAAAAAAGGGCG |
| ko | TGTCCCCAGATGGACAACTCCTC |
| | ACGAGGTGCTCCGTGTCC |
| adj | GCACAGACTGGCAAACTCAAAACAG |
| | ACCTGAAGTGCACATAATCCTGCC |
| CRISPR site 1a sgRNA | CACCTATAATCCGAAAGCAACCTC |
| | AAACGAGGTTGCTTTCGGATTATA |
| CRISPR site 1b sgRNA | CACCAGTCCTTTTACCTCTCTGGG |
| | AAACCCCAGAGAGGTAAAAGGACT |
| CRISPR site 2a sgRNA | CACCTATAATCCGGAGATAGCCTC |
| | AAACGAGGCTATCTCCGGATTATA |
| CRISPR site 2b sgRNA | CACCGAAGTTCAGCTCCTTTTCCA |
| | AAACTGGAAAAGGAGCTGAACTTC |
| CRISPR site 3a sgRNA | CACCTACAATCCGAAAGCAACCTC |
| | AAACGAGGTTGCTTTCGGATTGTA |
| CRISPR site 3b sgRNA | CACCAGCTCACTTTCCATAATAAA |
| | AAACTTTATTATGGAAAGTGAGCT |
| vtRNA1-1 probes | AGCGGTTACTTCGACAGTTCTTTAATTG |
| | AACAACCCAGACAGGTTGCTTG |
| vtRNA1-2 probes | AGCTCAGCGGTTACTTCGAGTAC |
| | CTGGAAAGCACCCGCGG |
| vtRNA1-3 probes | CAGCGGTTACTTCGCGTG |
| | ACAACCCAGAGAGGTGGTTTGATG |
| vtRNA2-1 probes | GCGAGACCGCATGACGC |
| | AGCCTCCATACTTTCTGTCACACC |
| SLFN11 probes | GTAGGGAAGTCCTGGGATGTGC |
| | GGGTTGGCAAAAATGAACACAAGG |
| GAPDH probes | GAGAAGGCTGGGGCTCATTT |
| | TGATGACCCTTTTGGCTCCC |
| GAPDH probes (Vir.) | GGAGCGAGATCCCTCCAAAAT |
| | GGCTGTTGTCATACTTCTCATGG |
| OC43 gRNA (RdRp) probes | GAGTGTAGATGCCCGTCTCG |
| | ATCAACACGCTGAAAACGGC |
| vtRNA1-1 template | AAAAGGACTGGAGAGCGCCCGCGGGTCTCGAACAACCCAGACAGGTTGCTTGTTTCAATTA<br>AAGAACTGTCGAAGTAACCGCTGAGCTAAAGCCAGCCTATAGTGAGTCGTATTA |
| vtRNA1-2 template | AAAAGAGCTGGAAAGCACCCGCGGGTCTCGAACCACCCAGAGAGGTGGTTACAATGTACT<br>CGAAGTAACCGCTGAGCTAAAGCCAGCCTATAGTGAGTCGTATTA |
| vtRNA1-3 template | AAGAGGGCTGGAGAGCGCCCGCGGGTCTCGAACAACCCAGAGAGGTGGTTTGATGACAC<br>GCGAAGTAACCGCTGAGCTAAAGCCAGCCTATAGTGAGTCGTATTA |
| vtRNA2-1 template | AAAAGGGTCAGTAAGCACCCGCGGGTCTCGAACCCCAGCACAGAGATGGACAGATAGAAA<br>GTCCGGCATGAGGAGGTAACCGCTTGAGCTAACTCCGACCTATAGTGAGTCGTATTA |
| T7 promoter | TAATACGACTCACTATA |
| VT11 insert | GATCTCTCACCAGTCAATAAAATATAATCCGAAAGCAACCTCCGGGCTGGCTTTAGCTCAGCGGTTACTT<br>CGACAGTTCTTTAATTGAAACAAGCAACCTGTCTGGGTTGTTCGAGACCCGCGGGCGCTCTCCAGTCCT<br>TTTACCTCTCTGGGCGGTTCGAGACCCGCGGGAGCTCTCCAGT |
| | AGCTACTGGAGAGCTCCCGCGGGTCTCGAACCGCCCAGAGAGGTAAAAGGACTGGAGAGCGCCCGCG<br>GGTCTCGAACAACCCAGACAGGTTGCTTGTTTCAATTAAAGAACTGTCGAAGTAACCGCTGAGCTAAAG<br>CCAGCCCGGAGGTTGCTTTCGGATTATATTTTATTGACTGGTGAGA |

| | |
|---|---|
| VT11del insert | GATCTCTCACCAGTCAATAAAATATAATCCGAAAGCTCCGGGCTGGCTTTAGCTCAGCGGTTACTTCGAC<br>AGTTCTTTAATTGAAACAAGCAACCTGTCTGGGTTGTTCGAGACCCGCGGGCGCTCTCCAGTCCTTTTA<br>CCTCTCTGGGCGGTTCGAGACCCGCGGGAGCTCTCCAGT |
| | AGCTACTGGAGAGCTCCCGCGGGTCTCGAACCGCCCAGAGAGGTAAAAGGACTGGAGAGCGCCCGCG<br>GGTCTCGAACAACCCAGACAGGTTGCTTGTTTCAATTAAAGAACTGTCGAAGTAACCGCTGAGCTAAAG<br>CCAGCCCGGAGCTTTCGGATTATATTTTATTGACTGGTGAGA |
| VT11 U6 insert | CACCGGCTGGCTTTAGCTCAGCGGTTACTTCGACAGTTCTTTAATTGAAACAAGCAACCTGT<br>CTGGGTTGTTCGAGACCCGCGGGCGCTCTCCAGTCCTTTT |
| | AAAAAAAAGGACTGGAGAGCGCCCGCGGGTCTCGAACAACCCAGACAGGTTGCTTGTTTC<br>AATTAAAGAACTGTCGAAGTAACCGCTGAGCTAAAGCCAGCC |
| VT21 insert | GATCCTCTTCTATGGTTTAGAAGTTTCAGTCGCACACTCCTACCCGGGTCGGAGTTAGCTCAAGCGGTTA<br>CCTCCTCATGCCGGACTTTCTATCTGTCCATCTCTGTGCTGGGGTTCGAGACCCGCGGGTGCTTACTGA<br>CCCTTTTATGCAATAAATTCGGTATAATCTGTCACTCTGAAGGCTTTGTTATT |
| | AGCTAATAACAAAGCCTTCAGAGTGACAGATTATACCGAATTTATTGCATAAAAGGGTCAGTAAGCACCCG<br>CGGGTCTCGAACCCCAGCACAGAGATGGACAGATAGAAAGTCCGGCATGAGGAGGTAACCGCTTGAGC<br>TAACTCCGACCCGGGTAGGAGTGTGCGACTGAAACTTCTAAACCATAGAAGAG |

**SUPPLEMENTAL TABLE 1. Synthetic oligos for these studies**. A table of all synthetic oligos used in these studies, presented 5'→3' in Integrated DNA Technologies (IDT; Coralville, Iowa USA) sequence format. The second set of GAPDH probes ("Vir.") correspond to those used in OC43-associated RT-qPCR experiments.

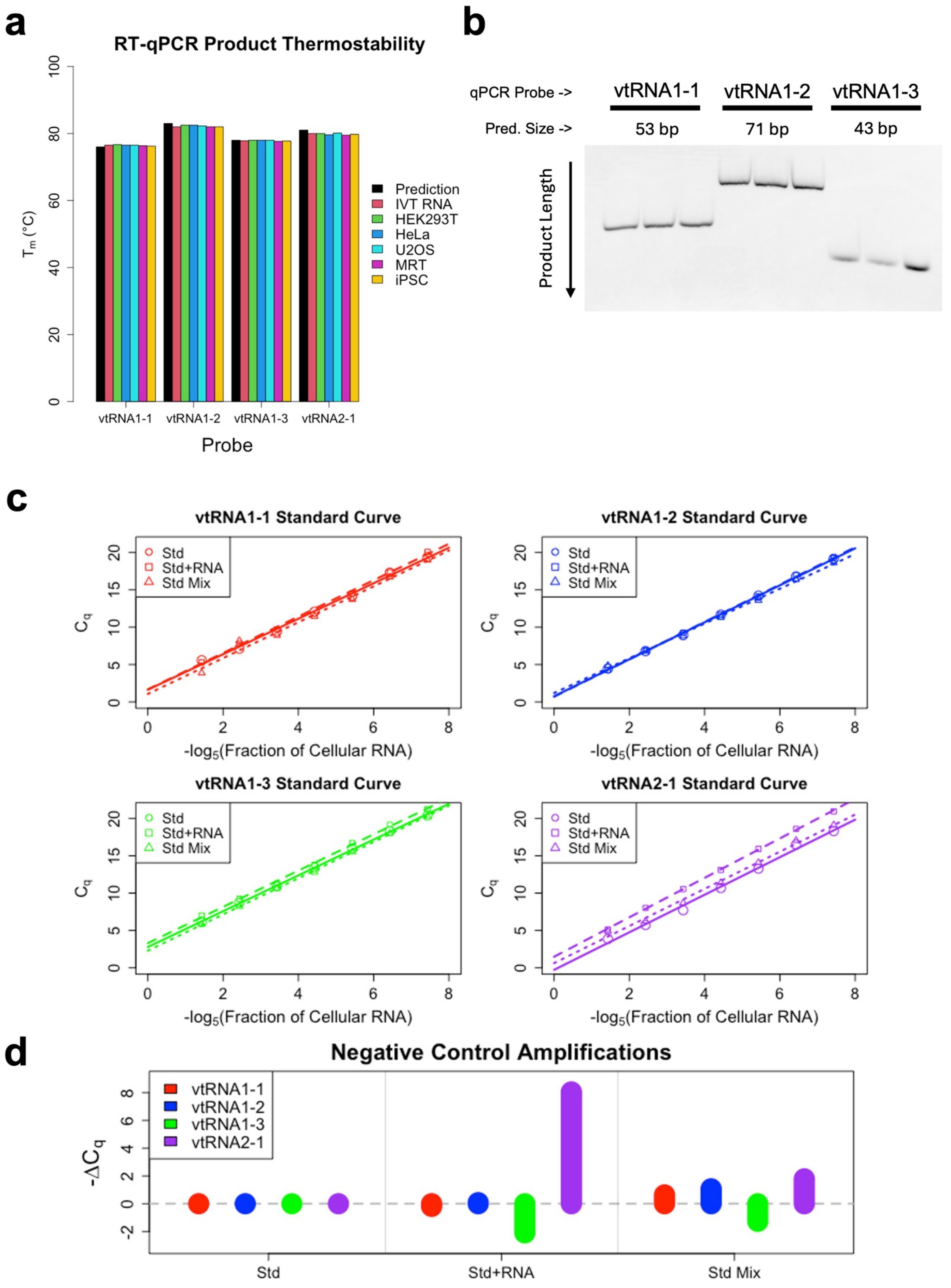
a
RT-qPCR Product Thermostability
Tm (°C)
Prediction
IVT RNA
HEK293T
HeLa
U2OS
MRT
iPSC
vtRNA1-1
vtRNA1-2
vtRNA1-3
vtRNA2-1
Probe
b
qPCR Probe ->
vtRNA1-1
vtRNA1-2
vtRNA1-3
Pred. Size ->
53 bp
71 bp
43 bp
Product Length
c
vtRNA1-1 Standard Curve
vtRNA1-2 Standard Curve
vtRNA1-3 Standard Curve
vtRNA2-1 Standard Curve
Std
Std+RNA
Std Mix
Cq
-log5(Fraction of Cellular RNA)
d
Negative Control Amplifications
-ΔCq
vtRNA1-1
vtRNA1-2
vtRNA1-3
vtRNA2-1
Std
Std+RNA
Std Mix

**SUPPLEMENTAL FIGURE 1. RT-qPCR probes are highly specific for vtRNA1 paralogs. [a]** Melt curves were generated for the amplification products from the Fig. 1a RT-qPCR experiment, and their melting temperatures ($T_m$) interpolated. “Prediction” refers to the theoretical $T_m$ for the anticipated qPCR product, calculated via the GeneGlobe Tm predictor (Qiagen). “IVT RNA” refers to the $T_m$ of samples from Fig. 1a standard curves. All other notation is consistent with Fig. 1a. Every data point is the average $T_m$ of at least n=2 samples from Fig. 1a. **[b]** The HEK293T samples’ final qPCR amplification products were visualized on a polyacrylamide gel to compare their expected lengths (“Pred. Size”) to relative migration (“Product Length”), and to assess their homogeneity. **[c]** Standard curves were determined by subjecting the vtRNA paralogs’ IVT products were subjected to RT-qPCR alone (“Std”, solid lines), in the presence of total RNA from the vt1-TKO c2 cell line (“Std+RNA”, dashed lines), or in a 4-paralog mix of the standard curves (“Std Mix”, dotted lines), then fit with linear regression. **[d]** Relative $C_q$ values of the zero-concentration samples in each vtRNA standard curve, normalized to the respective “Std” samples. The $-\Delta C_q$ values of the “Std+RNA” zero-concentration samples represent intrinsic vtRNA levels in the vt1-TKO c2 total RNA, while the “Std Mix” values are influenced by general variance.

*VTRNA1-1 Gene Locus:*

```
HEK293T_WT     ACTCTCTTTTTGACTCACTCTTTGTGACGTAGGTCTTTCTCACCAGTCAATAAAATATAA   60
CTRLdel        ACTCTCTTTTTGACTCACTCTTTGTGACGTAGGTCTTTCTCACCAGTCAATAAAATATAA   60
VTRNA1-1_KO    ACTCTCTTTTTGACTCACTCTTTGTGACGTAGGTCTTTCTCACCAGTCAATAAAATATAA   60
VTRNA1-2_KO    ACTCTCTTTTTGACTCACTCTTTGTGACGTAGGTCTTTCTCACCAGTCAATAAAATATAA   60
VTRNA1-3_KO    ACTCTCTTTTTGACTCACTCTTTGTGACGTAGGTCTTTCTCACCAGTCAATAAAATATAA   60
VTRNA1-TKOc1   ACTCTCTTTTTGACTCACTCTTTGTGACGTAGGTCTTTCTCACCAGTCAATAAAATATAA   60
VTRNA1-TKOc2   ACTCTCTTTTTGACTCACTCTTTGTGACGTAGGTCTTTCTCACCAGTCAATAAAATATAA   60
VTRNA1-TKOc3   ACTCTCTTTTTGACTCACTCTTTGTGACGTAGGTCTTTCTCACCAGTCAATAAAATATAA   60
HeLa           ACTCTCTTTTTGACTCACTCTTTGTGACGTAGGTCTTTCTCACCAGTCAATAAAATATAA   60
U2OS           ACTCTCTTTTTGACTCACTCTTTGTGACGTAGGTCTTTCTCACCAGTCAATAAAATATAA   60
MRT            ACTCTCTTTTTGACTCACTCTTTGTGACGTAGGTCTTTCTCACCAGTCAATAAAATATAA   60
BJ_Fib         ACTCTCTTTTTGACTCACTCTTTGTGACGTAGGTCTTTCTCACCAGTCAATAAAATATAA   60
iPSC           ACTCTCTTTTTGACTCACTCTTTGTGACGTAGGTCTTTCTCACCAGTCAATAAAATATAA   60

HEK293T_WT     TCCGAAAGCAACCTCCGGGCTGGCTTTAGCTCAGCGGTTACTTCGACAGTTCTTTAATTG   120
CTRLdel        TCCGAAAG----CTCCGGGCTGGCTTTAGCTCAGCGGTTACTTCGACAGTTCTTTAATTG   116
VTRNA1-1_KO    TCCGAAAGCAAC------------------------------------------------   72
VTRNA1-2_KO    TCCGAAAGCAACCTCCGGGCTGGCTTTAGCTCAGCGGTTACTTCGACAGTTCTTTAATTG   120
VTRNA1-3_KO    TCCGAAAGCAAC-TCCGGGCTGGCTTTAGCTCAGCGGTTACTTCGACAGTTCTTTAATTG   119
VTRNA1-TKOc1   TCCGAAAGCAAC------------------------------------------------   72
VTRNA1-TKOc2   TCCGAAAGCAAC------------------------------------------------   72
VTRNA1-TKOc3   TCCGAAAGCAAC------------------------------------------------   72
HeLa           TCCGAAAGCAACCTCCGGGCTGGCTTTAGCTCAGCGGTTACTTCGACAGTTCTTTAATTG   120
U2OS           TCCGAAAGCAACCTCCGGGCTGGCTTTAGCTCAGCGGTTACTTCGACAGTTCTTTAATTG   120
MRT            TCCGAAAGCAACCTCCGGGCTGGCTTTAGCTCAGCGGTTACTTCGACAGTTCTTTAATTG   120
BJ_Fib         TCCGAAAGCAACCTCCGGGCTGGCTTTAGCTCAGCGGTTACTTCGACAGTTCTTTAATTG   120
iPSC           TCCGAAAGCAACCTCCGGGCTGGCTTTAGCTCAGCGGTTACTTCGACAGTTCTTTAATTG   120

HEK293T_WT     AAACAAGCAACCTGTCTGGGTTGTTCGAGACCCGCGGGCGCTCTCCAGTCCTTTTACCTC   180
CTRLdel        AAACAAGCAACCTGTCTGGGTTGTTCGAGACCCGCGGGCGCTCTCCAGTCCTTTTACCTC   176
VTRNA1-1_KO    ------------------------------------------------------------   72
VTRNA1-2_KO    AAACAAGCAACCTGTCTGGGTTGTTCGAGACCCGCGGGCGCTCTCCAGTCCTTTTACCTC   180
```

```
VTRNA1-3_KO      AAACAAGCAACCTGTCTGGGTTGTTCGAGACCCGCGGGCGCTCTCCAGTCCTTTTACCTC     179
VTRNA1-TKOc1     ------------------------------------------------------------     72
VTRNA1-TKOc2     ------------------------------------------------------------     72
VTRNA1-TKOc3     ------------------------------------------------------------     72
HeLa             AAACAAGCAACCTGTCTGGGTTGTTCGAGACCCGCGGGCGCTCTCCAGTCCTTTTACCTC     180
U2OS             AAACAAGCAACCTGTCTGGGTTGTTCGAGACCCGCGGGCGCTCTCCAGTCCTTTTACCTC     180
MRT              AAACAAGCAACCTGTCTGGGTTGTTCGAGACCCGCGGGCGCTCTCCAGTCCTTTTACCTC     180
BJ_Fib           AAACAAGCAACCTGTCTGGGTTGTTCGAGACCCGCGGGCGCTCTCCAGTCCTTTTACCTC     180
iPSC             AAACAAGCAACCTGTCTGGGTTGTTCGAGACCCGCGGGCGCTCTCCAGTCCTTTTACCTC     180

HEK293T_WT       TCTGGGCGGTTCGAGACCCGCGGGAGCTCTCCAGTCCTTTTCACTGCTGAAGTTCAGCTC     240
CTRLdel          TCTGGGCGGTTCGAGACCCGCGGGAGCTCTCCAGTCCTTTTCACTGCTGAAGTTCAGCTC     236
VTRNA1-1_KO      --TGGGCGGTTCGAGACCCGCGGGAGCTCTCCAGTCCTTTTCACTGCTGAAGTTCAGCTC     130
VTRNA1-2_KO      TCTGGGCGGTTCGAGACCCGCGGGAGCTCTCCAGTCCTTTTCACTGCTGAAGTTCAGCTC     240
VTRNA1-3_KO      TCTGGGCGGTTCGAGACCCGCGGGAGCTCTCCAGTCCTTTTCACTGCTGAAGTTCAGCTC     239
VTRNA1-TKOc1     ------------------------------------------------------------     72
VTRNA1-TKOc2     ------------------------------------------------------------     72
VTRNA1-TKOc3     ------------------------------------------------------------     72
HeLa             TCTGGGCGGTTCGAGACCCGCGGGAGCTCTCCAGTCCTTTTCACTGCTGAAGTTCAGCTC     240
U2OS             TCTGGGCGGTTCGAGACCCGCGGGAGCTCTCCAGTCCTTTTCACTGCTGAAGTTCAGCTC     240
MRT              TCTGGGCGGTTCGAGACCCGCGGGAGCTCTCCAGTCCTTTTCACTGCTGAAGTTCAGCTC     240
BJ_Fib           TCTGGGCGGTTCGAGACCCGCGGGAGCTCTCCAGTCCTTTTCACTGCTGAAGTTCAGCTC     240
iPSC             TCTGGGCGGTTCGAGACCCGCGGGAGCTCTCCAGTCCTTTTCACTGCTGAAGTTCAGCTC     240

HEK293T_WT       CCTTTCCATGGTAAAAGGA     259
CTRLdel          CCTTTCCATGGTAAAAGGA     254
VTRNA1-1_KO      CCTTTCCATGGTAAAAGGA     148
VTRNA1-2_KO      CCTTTCCATGGTAAAAGGA     258
VTRNA1-3_KO      CCTTTCCATGGTAAAAGGA     257
VTRNA1-TKOc1     -------------------     71
VTRNA1-TKOc2     -------------------     71
VTRNA1-TKOc3     -------------------     71
HeLa             CCTTTCCATGGTAAAAGGA     259
U2OS             CCTTTCCATGGTAAAAGGA     259
```

```
MRT                CCTTTCCATGGTAAAAGGA     259
BJ_Fib             CCTTTCCATGGTAAAAGGA     259
iPSC               CCTTTCCATGGTAAAAGGA     259
```

*VTRNA1-2 Gene Locus:*

```
HEK293T_WT         ACTGTTGTTTTGACTCACTTTGTGACGTAGGTCTTTCTCACCAGTCAAGAAAATATAATC      60
CTRLdel            ACTGTTGTTTTGACTCACTTTGTGACGTAGGTCTTTCTCACCAGTCAAGAAAATATAATC      60
VTRNA1-1_KO        ACTGTTGTTTTGACTCACTTTGTGACGTAGGTCTTTCTCACCAGTCAAGAAAATATAATC      60
VTRNA1-2_KO        ACTGTTGTTTTGACTCACTTTGTGACGTAGGTCTTTCTCACCAGTCAAGAAAATATAATC      60
VTRNA1-3_KO        ACTGTTGTTTTGACTCACTTTGTGACGTAGGTCTTTCTCACCAGTCAAGAAAATATAATC      60
VTRNA1-TKOc1       ------------------------------------------------------------      0
VTRNA1-TKOc2       ------------------------------------------------------------      0
VTRNA1-TKOc3       ------------------------------------------------------------      0
HeLa               ACTGTTGTTTTGACTCACTTTGTGACGTAGGTCTTTCTCACCAGTCAAGAAAATATAATC      60
U2OS               ACTGTTGTTTTGACTCACTTTGTGACGTAGGTCTTTCTCACCAGTCAAGAAAATATAATC      60
MRT                ACTGTTGTTTTGACTCACTTTGTGACGTAGGTCTTTCTCACCAGTCAAGAAAATATAATC      60
BJ_Fib             ACTGTTGTTTTGACTCACTTTGTGACGTAGGTCTTTCTCACCAGTCAAGAAAATATAATC      60
iPSC               ACTGTTGTTTTGACTCACTTTGTGACGTAGGTCTTTCTCACCAGTCAAGAAAATATAATC      60

HEK293T_WT         CGGAGATAGCCTCTGGGCTGGCTTTAGCTCAGCGGTTACTTCGAGTACATTGTAACCACC      120
CTRLdel            CGGAGATAGCCTCTGGGCTGGCTTTAGCTCAGCGGTTACTTCGAGTACATTGTAACCACC      120
VTRNA1-1_KO        CGGAGATAGCCTCTGGGCTGGCTTTAGCTCAGCGGTTACTTCGAGTACATTGTAACCACC      120
VTRNA1-2_KO        CGGAGATAGC--------------------------------------------------      70
VTRNA1-3_KO        CGGAGATAGCCTCTGGGCTGGCTTTAGCTCAGCGGTTACTTCGAGTACATTGTAACCACC      120
VTRNA1-TKOc1       ------------------------------------------------------------      0
VTRNA1-TKOc2       ------------------------------------------------------------      0
VTRNA1-TKOc3       ------------------------------------------------------------      0
HeLa               CGGAGATAGCCTCTGGGCTGGCTTTAGCTCAGCGGTTACTTCGAGTACATTGTAACCACC      120
U2OS               CGGAGATAGCCTCTGGGCTGGCTTTAGCTCAGCGGTTACTTCGAGTACATTGTAACCACC      120
MRT                CGGAGATAGCCTCTGGGCTGGCTTTAGCTCAGCGGTTACTTCGAGTACATTGTAACCACC      120
BJ_Fib             CGGAGATAGCCTCTGGGCTGGCTTTAGCTCAGCGGTTACTTCGAGTACATTGTAACCACC      120
iPSC               CGGAGATAGCCTCTGGGCTGGCTTTAGCTCAGCGGTTACTTCGAGTACATTGTAACCACC      120
```

```
HEK293T_WT       TCTCTGGGTGGTTCGAGACCCGCGGGTGCTTTCCAGCTCTTTTACTGCTGAAGTTCAGCT      180
CTRLdel          TCTCTGGGTGGTTCGAGACCCGCGGGTGCTTTCCAGCTCTTTTACTGCTGAAGTTCAGCT      180
VTRNA1-1_KO      TCTCTGGGTGGTTCGAGACCCGCGGGTGCTTTCCAGCTCTTTTACTGCTGAAGTTCAGCT      180
VTRNA1-2_KO      ------------------------------------------------------------      70
VTRNA1-3_KO      TCTCTGGGTGGTTCGAGACCCGCGGGTGCTTTCCAGCTCTTTTACTGCTGAAGTTCAGCT      180
VTRNA1-TKOc1     ------------------------------------------------------------      0
VTRNA1-TKOc2     ------------------------------------------------------------      0
VTRNA1-TKOc3     ------------------------------------------------------------      0
HeLa             TCTCTGGGTGGTTCGAGACCCGCGGGTGCTTTCCAGCTCTTTTACTGCTGAAGTTCAGCT      180
U2OS             TCTCTGGGTGGTTCGAGACCCGCGGGTGCTTTCCAGCTCTTTTACTGCTGAAGTTCAGCT      180
MRT              TCTCTGGGTGGTTCGAGACCCGCGGGTGCTTTCCAGCTCTTTTACTGCTGAAGTTCAGCT      180
BJ_Fib           TCTCTGGGTGGTTCGAGACCCGCGGGTGCTTTCCAGCTCTTTTACTGCTGAAGTTCAGCT      180
iPSC             TCTCTGGGTGGTTCGAGACCCGCGGGTGCTTTCCAGCTCTTTTACTGCTGAAGTTCAGCT      180

HEK293T_WT       CCTTTTCCATGGGGAACCATGGAGAAATGGCTGATTTTAGCACTGGGCAGGAAATACAGA      240
CTRLdel          CCTTTTCCATGGGGAACCATGGAGAAATGGCTGATTTTAGCACTGGGCAGGAAATACAGA      240
VTRNA1-1_KO      CCTTTTCCATGGGGAACCATGGAGAAATGGCTGATTTTAGCACTGGGCAGGAAATACAGA      240
VTRNA1-2_KO      -----TCCATGGGGAACCATGGAGAAATGGCTGATTTTAGCACTGGGCAGGAAATACAGA      125
VTRNA1-3_KO      CCTTTTCCATGGGGAACCATGGAGAAATGGCTGATTTTAGCACTGGGCAGGAAATACAGA      240
VTRNA1-TKOc1     ------------------------------------------------------------      0
VTRNA1-TKOc2     ------------------------------------------------------------      0
VTRNA1-TKOc3     ------------------------------------------------------------      0
HeLa             CCTTTTCCATGGGGAACCATGGAGAAATGGCTGATTTTAGCACTGGGCAGGAAATACAGA      240
U2OS             CCTTTTCCATGGGGAACCATGGAGAAATGGCTGATTTTAGCACTGGGCAGGAAATACAGA      240
MRT              CCTTTTCCATGGGGAACCATGGAGAAATGGCTGATTTTAGCACTGGGCAGGAAATACAGA      240
BJ_Fib           CCTTTTCCATGGGGAACCATGGAGAAATGGCTGATTTTAGCACTGGGCAGGAAATACAGA      240
iPSC             CCTTTTCCATGGGGAACCATGGAGAAATGGCTGATTTTAGCACTGGGCAGGAAATACAGA      240

HEK293T_WT       G     241
CTRLdel          G     241
VTRNA1-1_KO      G     241
VTRNA1-2_KO      G     126
VTRNA1-3_KO      G     241
```

```
VTRNA1-TKOc1      -     0
VTRNA1-TKOc2      -     0
VTRNA1-TKOc3      -     0
HeLa              G     241
U2OS              G     241
MRT               G     241
BJ_Fib            G     241
iPSC              G     241
```

*VTRNA1-3 Gene Locus:*

```
HEK293T_WT        CACCGTTGTTTTGACTCACTCTTTGTGACGTAGGTCTTTCTCACCAGTCAAGAAAATACA      60
CTRLdel           CACCGTTGTTTTGACTCACTCTTTGTGACGTAGGTCTTTCTCACCAGTCAAGAAAATACA      60
VTRNA1-1_KO       CACCGTTGTTTTGACTCACTCTTTGTGACGTAGGTCTTTCTCACCAGTCAAGAAAATACA      60
VTRNA1-2_KO       CACCGTTGTTTTGACTCACTCTTTGTGACGTAGGTCTTTCTCACCAGTCAAGAAAATACA      60
VTRNA1-3_KO       CACCGTTGTTTTGACTCACTCTTTGTGACGTAGGTCTTTCTCACCAGTCAAGAAAATACA      60
VTRNA1-TKOc1      ------------------------------------------------------------      0
VTRNA1-TKOc2      ------------------------------------------------------------      0
VTRNA1-TKOc3      ------------------------------------------------------------      0
HeLa              CACCGTTGTTTTGACTCACTCTTTGTGACGTAGGTCTTTCTCACCAGTCAAGAAAATACA      60
U2OS              CACCGTTGTTTTGACTCACTCTTTGTGACGTAGGTCTTTCTCACCAGTCAAGAAAATACA      60
MRT               CACCGTTGTTTTGACTCACTCTTTGTGACGTAGGTCTTTCTCACCAGTCAAGAAAATACA      60
BJ_Fib            CACCGTTGTTTTGACTCACTCTTTGTGACGTAGGTCTTTCTCACCAGTCAAGAAAATACA      60
iPSC              CACCGTTGTTTTGACTCACTCTTTGTGACGTAGGTCTTTCTCACCAGTCAAGAAAATACA      60

HEK293T_WT        ATCCGAAAGCAACCTCTGGGCTGGCTTTAGCTCAGCGGTTACTTCGCGTGTCATCAAACC      120
CTRLdel           ATCCGAAAG----CTCTGGGCTGGCTTTAGCTCAGCGGTTACTTCGCGTGTCATCAAACC      116
VTRNA1-1_KO       ATCCGAAAGCAACCTCTGGGCTGGCTTTAGCTCAGCGGTTACTTCGCGTGTCATCAAACC      120
VTRNA1-2_KO       ATCCGAAAGCAACCTCTGGGCTGGCTTTAGCTCAGCGGTTACTTCGCGTGTCATCAAACC      120
VTRNA1-3_KO       ATCCGAAAGCAAC-----------------------------------------------      73
VTRNA1-TKOc1      ------------------------------------------------------------      0
VTRNA1-TKOc2      ------------------------------------------------------------      0
VTRNA1-TKOc3      ------------------------------------------------------------      0
```

```
HeLa            ATCCGAAAGCAACCTCTGGGCTGGCTTTAGCTCAGCGGTTACTTCGCGTGTCATCAAACC    120
U2OS            ATCCGAAAGCAACCTCTGGGCTGGCTTTAGCTCAGCGGTTACTTCGCGTGTCATCAAACC    120
MRT             ATCCGAAAGCAACCTCTGGGCTGGCTTTAGCTCAGCGGTTACTTCGCGTGTCATCAAACC    120
BJ_Fib          ATCCGAAAGCAACCTCTGGGCTGGCTTTAGCTCAGCGGTTACTTCGCGTGTCATCAAACC    120
iPSC            ATCCGAAAGCAACCTCTGGGCTGGCTTTAGCTCAGCGGTTACTTCGCGTGTCATCAAACC    120

HEK293T_WT      ACCTCTCTGGGTTGTTCGAGACCCGCGGGCGCTCTCCAGCCCTCTTACTGCTGAAGTTCA    180
CTRLdel         ACCTCTCTGGGTTGTTCGAGACCCGCGGGCGCTCTCCAGCCCTCTTACTGCTGAAGTTCA    176
VTRNA1-1_KO     ACCTCTCTGGGTTGTTCGAGACCCGCGGGCGCTCTCCAGCCCTCTTACTGCTGAAGTTCA    180
VTRNA1-2_KO     ACCTCTCTGGGTTGTTCGAGACCCGCGGGCGCTCTCCAGCCCTCTTACTGCTGAAGTTCA    180
VTRNA1-3_KO     ------------------------------------------------------------    73
VTRNA1-TKOc1    ------------------------------------------------------------    0
VTRNA1-TKOc2    ------------------------------------------------------------    0
VTRNA1-TKOc3    ------------------------------------------------------------    0
HeLa            ACCTCTCTGGGTTGTTCGAGACCCGCGGGCGCTCTCCAGCCCTCTTACTGCTGAAGTTCA    180
U2OS            ACCTCTCTGGGTTGTTCGAGACCCGCGGGCGCTCTCCAGCCCTCTTACTGCTGAAGTTCA    180
MRT             ACCTCTCTGGGTTGTTCGAGACCCGCGGGCGCTCTCCAGCCCTCTTACTGCTGAAGTTCA    180
BJ_Fib          ACCTCTCTGGGTTGTTCGAGACCCGCGGGCGCTCTCCAGCCCTCTTACTGCTGAAGTTCA    180
iPSC            ACCTCTCTGGGTTGTTCGAGACCCGCGGGCGCTCTCCAGCCCTCTTACTGCTGAAGTTCA    180

HEK293T_WT      GCTCACTTTCCATAATAAAAGGATCCAGGGTTTCTTGGAGAAATGGCTGATTTTAGTAGT    240
CTRLdel         GCTCACTTTCCAT--TAAAAGGATCCAGGGTTTCTTGGAGAAATGGCTGATTTTAGTAGT    234
VTRNA1-1_KO     GCTCACTTTCCATAATAAAAGGATCCAGGGTTTCTTGGAGAAATGGCTGATTTTAGTAGT    240
VTRNA1-2_KO     GCTCACTTTCCATAATAAAAGGATCCAGGGTTTCTTGGAGAAATGGCTGATTTTAGTAGT    240
VTRNA1-3_KO     ---------------TAAAAGGATCCAGGGTTTCTTGGAGAAATGGCTGATTTTAGTAGT    118
VTRNA1-TKOc1    -----------------AAAAGGATCCAGGGTTTCTTGGAGAAATGGCTGATTTTAGTAGT    44
VTRNA1-TKOc2    -----------------AAAAGGATCCAGGGTTTCTTGGAGAAATGGCTGATTTTAGTAGT    44
VTRNA1-TKOc3    ---------------TAAAAGGATCCAGGGTTTCTTGGAGAAATGGCTGATTTTAGTAGT    45
HeLa            GCTCACTTTCCATAATAAAAGGATCCAGGGTTTCTTGGAGAAATGGCTGATTTTAGTAGT    240
U2OS            GCTCACTTTCCATAATAAAAGGATCCAGGGTTTCTTGGAGAAATGGCTGATTTTAGTAGT    240
MRT             GCTCACTTTCCATAATAAAAGGATCCAGGGTTTCTTGGAGAAATGGCTGATTTTAGTAGT    240
BJ_Fib          GCTCACTTTCCATAATAAAAGGATCCAGGGTTTCTTGGAGAAATGGCTGATTTTAGTAGT    240
iPSC            GCTCACTTTCCATAATAAAAGGATCCAGGGTTTCTTGGAGAAATGGCTGATTTTAGTAGT    240
```

| | | |
|---|---|---|
| HEK293T_WT | GGGCAG | 246 |
| CTRLdel | GGGCAG | 240 |
| VTRNA1-1_KO | GGGCAG | 246 |
| VTRNA1-2_KO | GGGCAG | 246 |
| VTRNA1-3_KO | GGGCAG | 124 |
| VTRNA1-TKOc1 | GGGCAG | 50 |
| VTRNA1-TKOc2 | GGGCAG | 50 |
| VTRNA1-TKOc3 | GGGCAG | 51 |
| HeLa | GGGCAG | 246 |
| U2OS | GGGCAG | 246 |
| MRT | GGGCAG | 246 |
| BJ_Fib | GGGCAG | 246 |
| iPSC | GGGCAG | 246 |

*VTRNA2-1 Gene Locus:*

| | | |
|---|---|---|
| HEK293T_WT | GCGAGACCGCATGACGCAGGCCTCTCGGGGGCGGGGAGCACATGCAGCCCGTCCCTCTCC | 60 |
| CTRLdel | GCGAGACCGCATGACGCAGGCCTCTCGGGGGCGGGGAGCACATGCAGCCCGTCCCTCTCC | 60 |
| VTRNA1-1_KO | GCGAGACCGCATGACGCAGGCCTCTCGGGGGCGGGGAGCACATGCAGCCCGTCCCTCTCC | 60 |
| VTRNA1-2_KO | GCGAGACCGCATGACGCAGGCCTCTCGGGGGCGGGGAGCACATGCAGCCCGTCCCTCTCC | 60 |
| VTRNA1-3_KO | GCGAGACCGCATGACGCAGGCCTCTCGGGGGCGGGGAGCACATGCAGCCCGTCCCTCTCC | 60 |
| VTRNA1-TKOc1 | GCGAGACCGCATGACGCAGGCCTCTCGGGGGCGGGGAGCACATGCAGCCCGTCCCTCTCC | 60 |
| VTRNA1-TKOc2 | GCGAGACCGCATGACGCAGGCCTCTCGGGGGCGGGGAGCACATGCAGCCCGTCCCTCTCC | 60 |
| VTRNA1-TKOc3 | GCGAGACCGCATGACGCAGGCCTCTCGGGGGCGGGGAGCACATGCAGCCCGTCCCTCTCC | 60 |
| HeLa | GCGAGACCGCATGACGCAGGCCTCTCGGGGGCGGGGAGCACATGCAGCCCGTCCCTCTCC | 60 |
| U2OS | GCGAGACCGCATGACGCAGGCCTCTCGGGGGCGGGGAGCACATGCAGCCCGTCCCTCTCC | 60 |
| MRT | GCGAGACCGCATGACGCAGGCCTCTCGGGGGCGGGGAGCACATGCAGCCCGTCCCTCTCC | 60 |
| BJ_Fib | GCGAGACCGCATGACGCAGGCCTCTCGGGGGCGGGGAGCACATGCAGCCCGTCCCTCTCC | 60 |
| iPSC | GCGAGACCGCATGACGCAGGCCTCTCGGGGGCGGGGAGCACATGCAGCCCGTCCCTCTCC | 60 |

| | | |
|---|---|---|
| HEK293T_WT | ACATCGTCACTCTTCTATGGTTTAGAAGTTTCAGTCGCACACTCCTACCCGGGTCGGAGT | 120 |
| CTRLdel | ACATCGTCACTCTTCTATGGTTTAGAAGTTTCAGTCGCACACTCCTACCCGGGTCGGAGT | 120 |
| VTRNA1-1_KO | ACATCGTCACTCTTCTATGGTTTAGAAGTTTCAGTCGCACACTCCTACCCGGGTCGGAGT | 120 |

```
VTRNA1-2_KO        ACATCGTCACTCTTCTATGGTTTAGAAGTTTCAGTCGCACACTCCTACCCGGGTCGGAGT    120
VTRNA1-3_KO        ACATCGTCACTCTTCTATGGTTTAGAAGTTTCAGTCGCACACTCCTACCCGGGTCGGAGT    120
VTRNA1-TKOc1       ACATCGTCACTCTTCTATGGTTTAGAAGTTTCAGTCGCACACTCCTACCCGGGTCGGAGT    120
VTRNA1-TKOc2       ACATCGTCACTCTTCTATGGTTTAGAAGTTTCAGTCGCACACTCCTACCCGGGTCGGAGT    120
VTRNA1-TKOc3       ACATCGTCACTCTTCTATGGTTTAGAAGTTTCAGTCGCACACTCCTACCCGGGTCGGAGT    120
HeLa               ACATCGTCACTCTTCTATGGTTTAGAAGTTTCAGTCGCACACTCCTACCCGGGTCGGAGT    120
U2OS               ACATCGTCACTCTTCTATGGTTTAGAAGTTTCAGTCGCACACTCCTACCCGGGTCGGAGT    120
MRT                ACATCGTCACTCTTCTATGGTTTAGAAGTTTCAGTCGCACACTCCTACCCGGGTCGGAGT    120
BJ_Fib             ACATCGTCACTCTTCTATGGTTTAGAAGTTTCAGTCGCACACTCCTACCCGGGTCGGAGT    120
iPSC               ACATCGTCACTCTTCTATGGTTTAGAAGTTTCAGTCGCACACTCCTACCCGGGTCGGAGT    120

HEK293T_WT         TAGCTCAAGCGGTTACCTCCTCATGCCGGACTTTCTATCTGTCCATCTCTGTGCTGGGGT    180
CTRLdel            TAGCTCAAGCGGTTACCTCCTCATGCCGGACTTTCTATCTGTCCATCTCTGTGCTGGGGT    180
VTRNA1-1_KO        TAGCTCAAGCGGTTACCTCCTCATGCCGGACTTTCTATCTGTCCATCTCTGTGCTGGGGT    180
VTRNA1-2_KO        TAGCTCAAGCGGTTACCTCCTCATGCCGGACTTTCTATCTGTCCATCTCTGTGCTGGGGT    180
VTRNA1-3_KO        TAGCTCAAGCGGTTACCTCCTCATGCCGGACTTTCTATCTGTCCATCTCTGTGCTGGGGT    180
VTRNA1-TKOc1       TAGCTCAAGCGGTTACCTCCTCATGCCGGACTTTCTATCTGTCCATCTCTGTGCTGGGGT    180
VTRNA1-TKOc2       TAGCTCAAGCGGTTACCTCCTCATGCCGGACTTTCTATCTGTCCATCTCTGTGCTGGGGT    180
VTRNA1-TKOc3       TAGCTCAAGCGGTTACCTCCTCATGCCGGACTTTCTATCTGTCCATCTCTGTGCTGGGGT    180
HeLa               TAGCTCAAGCGGTTACCTCCTCATGCCGGACTTTCTATCTGTCCATCTCTGTGCTGGGGT    180
U2OS               TAGCTCAAGCGGTTACCTCCTCATGCCGGACTTTCTATCTGTCCATCTCTGTGCTGGGGT    180
MRT                TAGCTCAAGCGGTTACCTCCTCATGCCGGACTTTCTATCTGTCCATCTCTGTGCTGGGGT    180
BJ_Fib             TAGCTCAAGCGGTTACCTCCTCATGCCGGACTTTCTATCTGTCCATCTCTGTGCTGGGGT    180
iPSC               TAGCTCAAGCGGTTACCTCCTCATGCCGGACTTTCTATCTGTCCATCTCTGTGCTGGGGT    180

HEK293T_WT         TCGAGACCCGCGGGTGCTTACTGACCCTTTTATGCAATAAATTCGGTATAATCTGTCACT    240
CTRLdel            TCGAGACCCGCGGGTGCTTACTGACCCTTTTATGCAATAAATTCGGTATAATCTGTCACT    240
VTRNA1-1_KO        TCGAGACCCGCGGGTGCTTACTGACCCTTTTATGCAATAAATTCGGTATAATCTGTCACT    240
VTRNA1-2_KO        TCGAGACCCGCGGGTGCTTACTGACCCTTTTATGCAATAAATTCGGTATAATCTGTCACT    240
VTRNA1-3_KO        TCGAGACCCGCGGGTGCTTACTGACCCTTTTATGCAATAAATTCGGTATAATCTGTCACT    240
VTRNA1-TKOc1       TCGAGACCCGCGGGTGCTTACTGACCCTTTTATGCAATAAATTCGGTATAATCTGTCACT    240
VTRNA1-TKOc2       TCGAGACCCGCGGGTGCTTACTGACCCTTTTATGCAATAAATTCGGTATAATCTGTCACT    240
VTRNA1-TKOc3       TCGAGACCCGCGGGTGCTTACTGACCCTTTTATGCAATAAATTCGGTATAATCTGTCACT    240
HeLa               TCGAGACCCGCGGGTGCTTACTGACCCTTTTATGCAATAAATTCGGTATAATCTGTCACT    240
```

```
U2OS            TCGAGACCCGCGGGTGCTTACTGACCCTTTTATGCAATAAATTCGGTATAATCTGTCACT    240
MRT             TCGAGACCCGCGGGTGCTTACTGACCCTTTTATGCAATAAATTCGGTATAATCTGTCACT    240
BJ_Fib          TCGAGACCCGCGGGTGCTTACTGACCCTTTTATGCAATAAATTCGGTATAATCTGTCACT    240
iPSC            TCGAGACCCGCGGGTGCTTACTGACCCTTTTATGCAATAAATTCGGTATAATCTGTCACT    240

HEK293T_WT      CTGAAGGCTTTGTTATTTTTTATCCCTTTTAACTTTGCTAAATTAATAGGCTGATAACAT    300
CTRLdel         CTGAAGGCTTTGTTATTTTTTATCCCTTTTAACTTTGCTAAATTAATAGGCTGATAACAT    300
VTRNA1-1_KO     CTGAAGGCTTTGTTATTTTTTATCCCTTTTAACTTTGCTAAATTAATAGGCTGATAACAT    300
VTRNA1-2_KO     CTGAAGGCTTTGTTATTTTTTATCCCTTTTAACTTTGCTAAATTAATAGGCTGATAACAT    300
VTRNA1-3_KO     CTGAAGGCTTTGTTATTTTTTATCCCTTTTAACTTTGCTAAATTAATAGGCTGATAACAT    300
VTRNA1-TKOc1    CTGAAGGCTTTGTTATTTTTTATCCCTTTTAACTTTGCTAAATTAATAGGCTGATAACAT    300
VTRNA1-TKOc2    CTGAAGGCTTTGTTATTTTTTATCCCTTTTAACTTTGCTAAATTAATAGGCTGATAACAT    300
VTRNA1-TKOc3    CTGAAGGCTTTGTTATTTTTTATCCCTTTTAACTTTGCTAAATTAATAGGCTGATAACAT    300
HeLa            CTGAAGGCTTTGTTATTTTTTATCCCTTTTAACTTTGCTAAATTAATAGGCTGATAACAT    300
U2OS            CTGAAGGCTTTGTTATTTTTTATCCCTTTTAACTTTGCTAAATTAATAGGCTGATAACAT    300
MRT             CTGAAGGCTTTGTTATTTTTTATCCCTTTTAACTTTGCTAAATTAATAGGCTGATAACAT    300
BJ_Fib          CTGAAGGCTTTGTTATTTTTTATCCCTTTTAACTTTGCTAAATTAATAGGCTGATAACAT    300
iPSC            CTGAAGGCTTTGTTATTTTTTATCCCTTTTAACTTTGCTAAATTAATAGGCTGATAACAT    300

HEK293T_WT      AAGTTGGTGTCACTTTGAAGGTGTGACAGAAAGTATGGAGGCT     343
CTRLdel         AAGTTGGTGTCACTTTGAAGGTGTGACAGAAAGTATGGAGGCT     343
VTRNA1-1_KO     AAGTTGGTGTCACTTTGAAGGTGTGACAGAAAGTATGGAGGCT     343
VTRNA1-2_KO     AAGTTGGTGTCACTTTGAAGGTGTGACAGAAAGTATGGAGGCT     343
VTRNA1-3_KO     AAGTTGGTGTCACTTTGAAGGTGTGACAGAAAGTATGGAGGCT     343
VTRNA1-TKOc1    AAGTTGGTGTCACTTTGAAGGTGTGACAGAAAGTATGGAGGCT     343
VTRNA1-TKOc2    AAGTTGGTGTCACTTTGAAGGTGTGACAGAAAGTATGGAGGCT     343
VTRNA1-TKOc3    AAGTTGGTGTCACTTTGAAGGTGTGACAGAAAGTATGGAGGCT     343
HeLa            AAGTTGGTGTCACTTTGAAGGTGTGACAGAAAGTATGGAGGCT     343
U2OS            AAGTTGGTGTCACTTTGAAGGTGTGACAGAAAGTATGGAGGCT     343
MRT             AAGTTGGTGTCACTTTGAAGGTGTGACAGAAAGTATGGAGGCT     343
BJ_Fib          AAGTTGGTGTCACTTTGAAGGTGTGACAGAAAGTATGGAGGCT     343
iPSC            AAGTTGGTGTCACTTTGAAGGTGTGACAGAAAGTATGGAGGCT     343
```

**SUPPLEMENTAL FIGURE 2. Alignment of Nanopore sequencing results for VTRNA1 and 2 loci of various human cell lines.** Wild-type, CTRLdel, and vtRNA single-knockout cell lines were subjected to PCR with the vt11, vt12, vt13, and vt21 primer sets (Fig. 2a), and vtRNA triple-knockout cell lines were subjected to PCR with the ko and vt21 primer sets (Fig. 2a). Products were Nanopore sequenced and dominant sequences were aligned across the four vtRNA paralogs' genetic loci with the Clustal Omega Multiple Sequence Alignment (EMBL-EBI). Cas9 cleavage sites are highlighted in purple, vtRNA coding strand sequences in green, and transcription start and termination sites in blue and red, respectively.

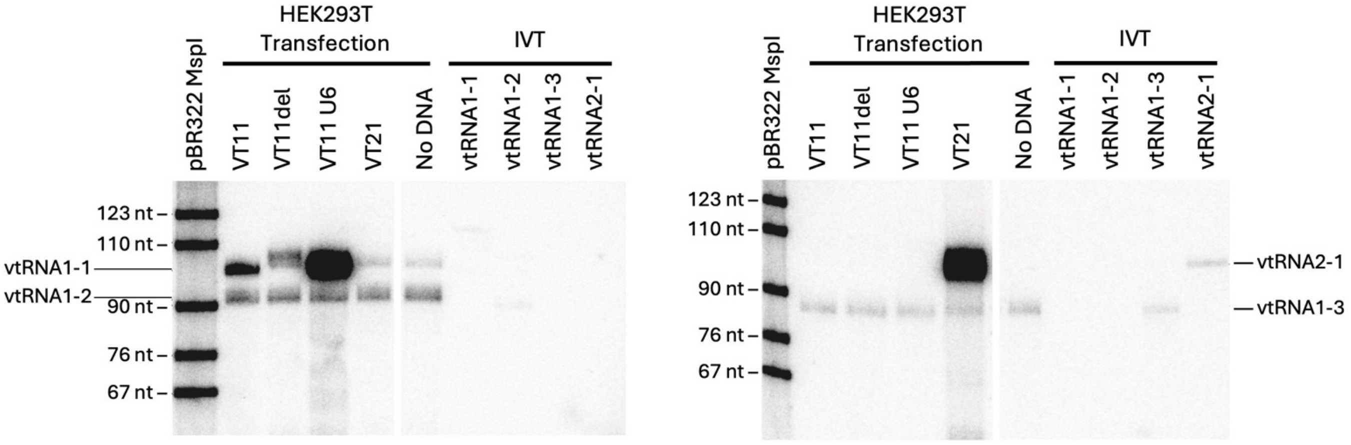


**SUPPLEMENTAL FIGURE 3. vtRNA2-1 expression in HEK293T cells is suppressed epigenetically.** Northern blots probing for vtRNA1-1 and 1-2 (left), and vtRNA1-3 and vtRNA2-1 (right) in cellular RNA extracted from wild-type HEK293T cells transfected with vtRNA-expressing plasmids. Constructs expressed vtRNAs using their natural, internal promoters (pUC19 vector + gene with 40 bp flanking sequences); that is, VT11 is $N_{40}$[VTRNA1-1]$N_{40}$, VT11del is $N_{31}\Delta_4 N_5$[VTRNA1-1]$N_{40}$, and VT21 is $N_{40}$[VTRNA2-1]$N_{40}$. A no-transfection control (No DNA) and a positive control with *VTRNA1-1* behind a synthetic U6 promoter (VT11 U6) are included. The negative (no-transfection) control shows the basal levels of vtRNAs in the wild-type cells. “HEK293T Transfection” lanes are loaded with equal amounts of cellular RNA. IVT RNA standards (“IVT”) are included, and the anticipated vtRNA product lengths are indicated by labels to the left/right of the gels. White, dividing lines in gels are from trimming extraneous lanes; images are cropped portions of the same parent image, without alterations to contrast, alignment, etc. Data are representative of three independent experiments.

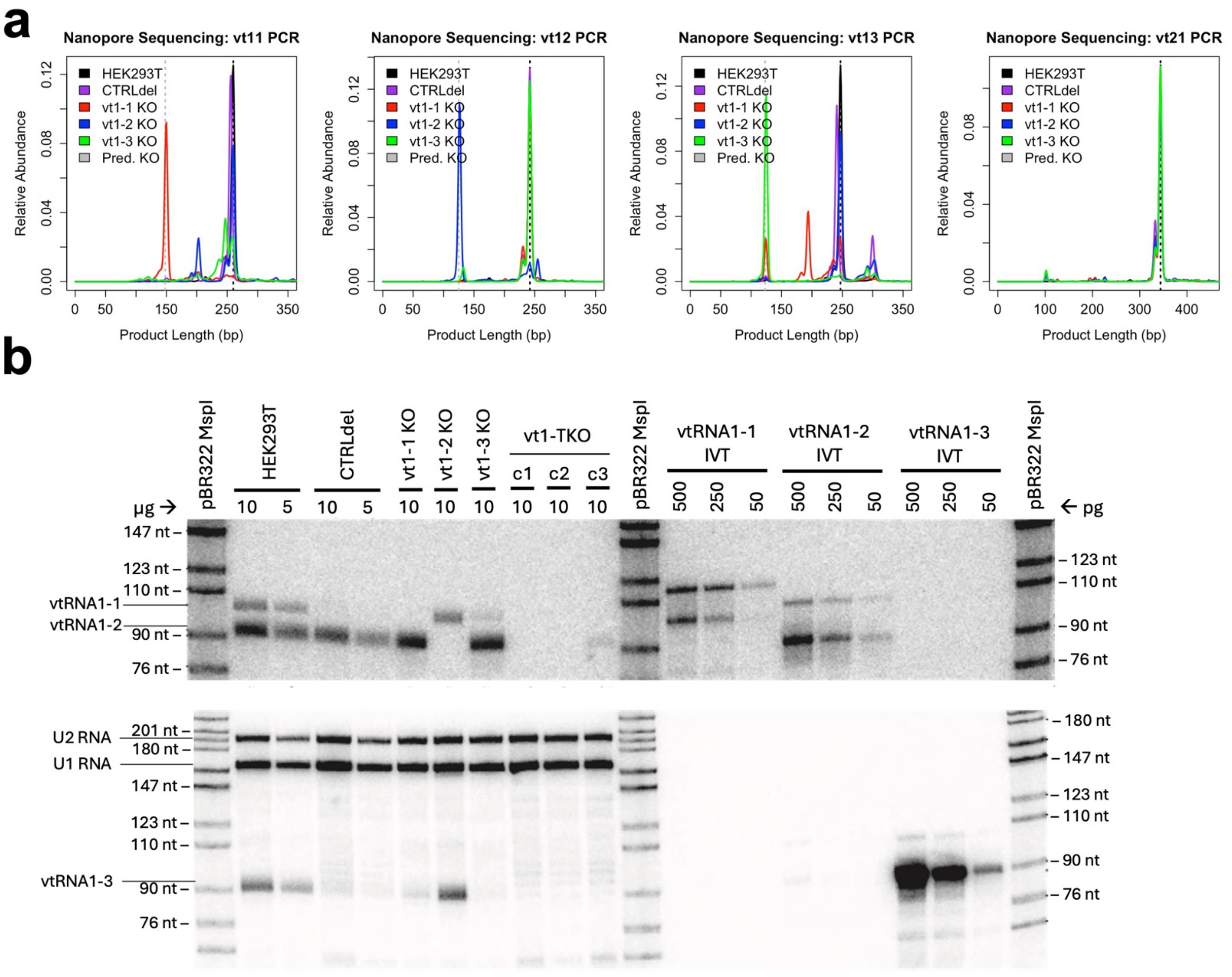


**SUPPLEMENTAL FIGURE 4. Validation of vtRNA1-1, 1-2, 1-3, and 1-TKO HEK293T cell lines.** [a] Product size distributions from Nanopore sequencing of Fig. 2b PCR reactions. Predicted product lengths for knockout cell lines ("Pred. KO"; vertical, grey dashed lines) are based on the Cas9 cleavage sites shown in Supplemental Figure 2. Wild-type length products (vertical, black dashed lines) are based on the dominant band in HEK293T cell line. [b] Northern blots probing for vtRNA1-1 and 1-2 (top), and vtRNA1-3 (bottom) in wild-type, CTRLdel, and vtRNA knockout HEK293T cell lines, and in IVT RNA standards ("IVT").

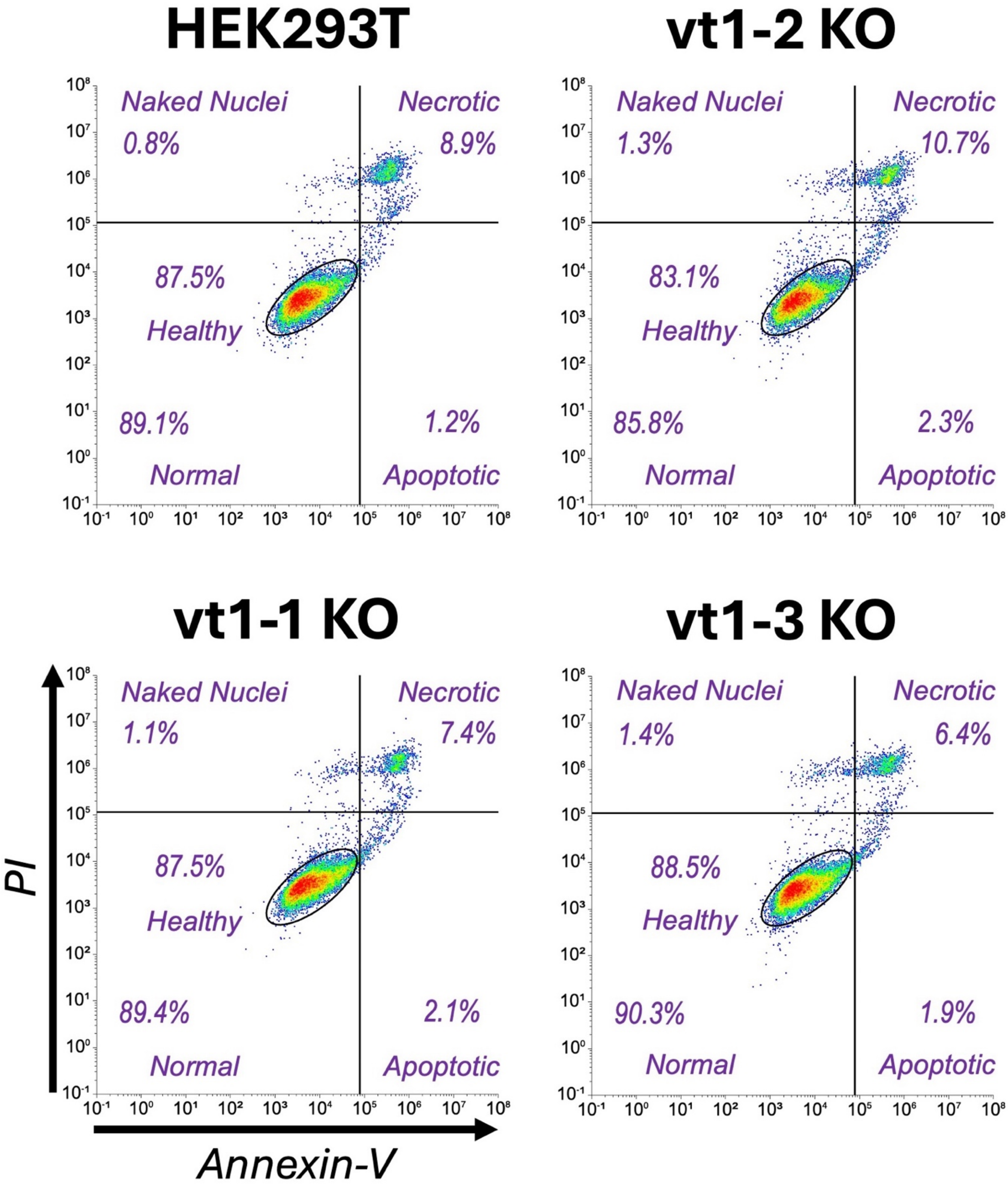


**SUPPLEMENTAL FIGURE 5. Cell viability of HEK293T lacking vtRNA1 paralogs is not compromised.** Wild-type or vtRNA1 single-knockout HEK293T cells were grown under standard conditions, dual stained with propidium iodide (PI) and Annexin-V, then analyzed via flow cytometry. Events shown were first gated by SSC-A/FSC-A then FSC-A/FSC-H to identify singlets. Gating and graphics were via the Floreada web server.

**a**

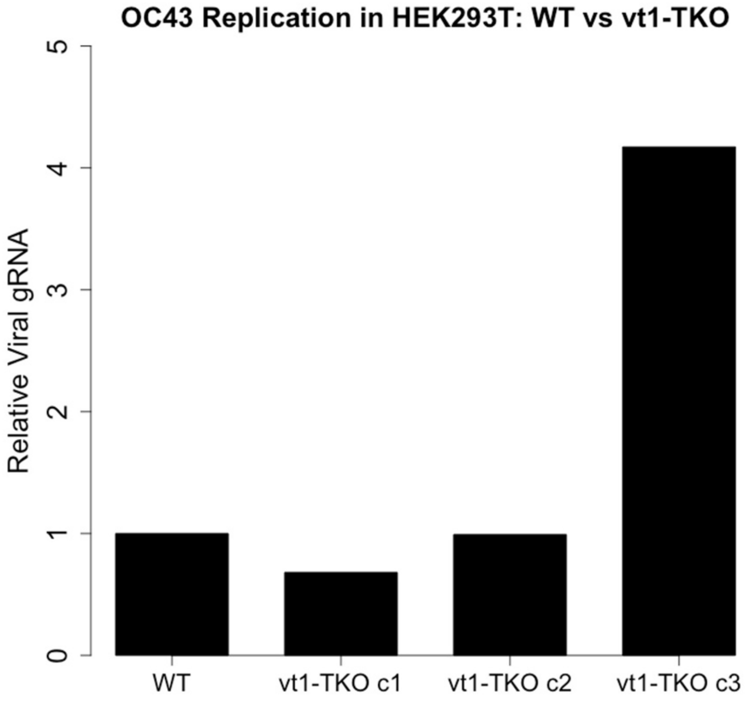
OC43 Replication in HEK293T: WT vs vt1-TKO
Relative Viral gRNA
0
1
2
3
4
5
WT
vt1-TKO c1
vt1-TKO c2
vt1-TKO c3

**b**

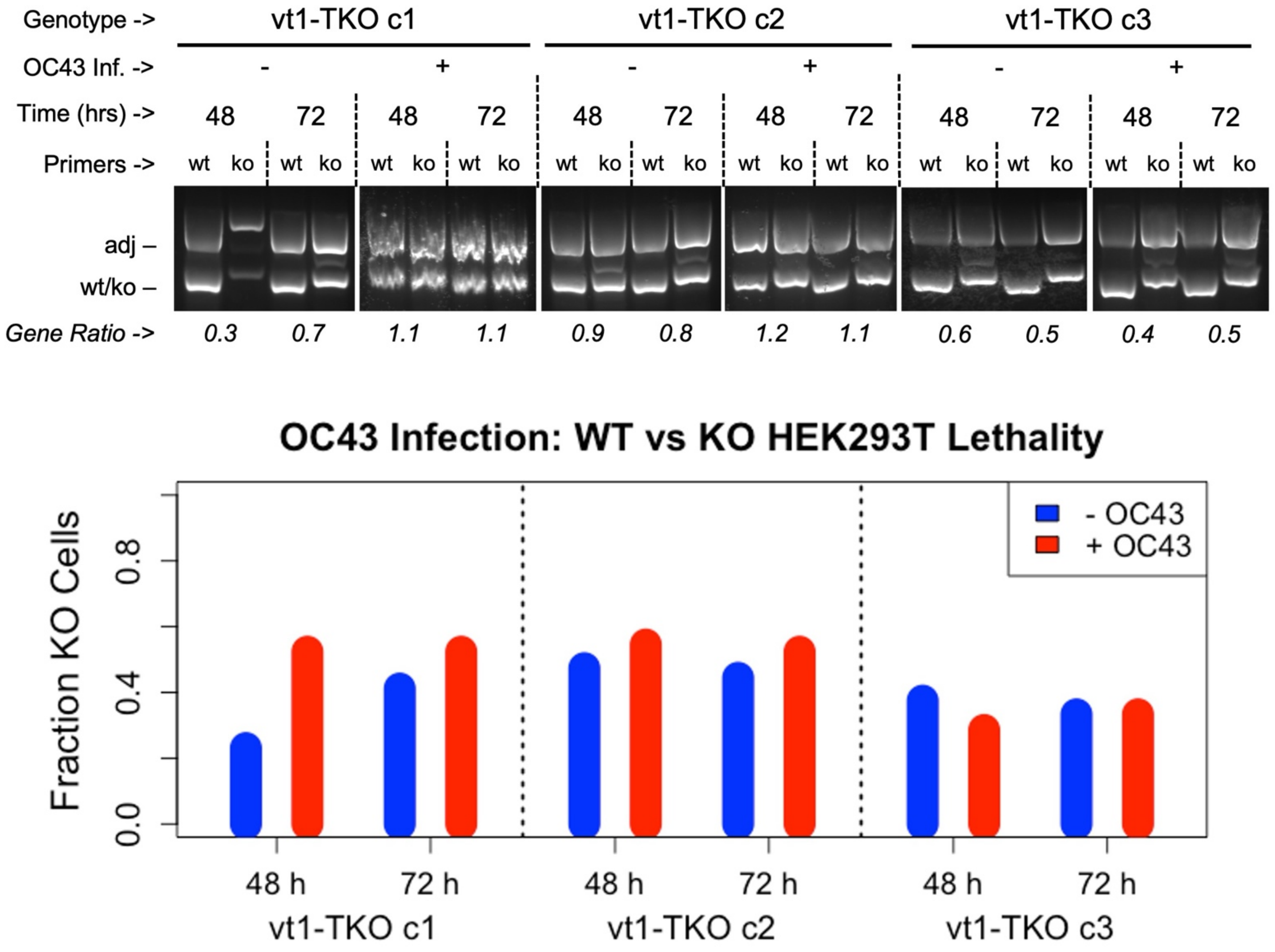
Genotype ->
vt1-TKO c1
vt1-TKO c2
vt1-TKO c3
OC43 Inf. ->
-
+
Time (hrs) ->
48
72
Primers ->
wt
ko
adj –
wt/ko –
Gene Ratio ->
0.3 0.7 1.1 1.1 0.9 0.8 1.2 1.1 0.6 0.5 0.4 0.5
OC43 Infection: WT vs KO HEK293T Lethality
Fraction KO Cells
0.0
0.4
0.8
- OC43
+ OC43
48 h
72 h
vt1-TKO c1
vt1-TKO c2
vt1-TKO c3

**SUPPLEMENTAL FIGURE 6. Loss of vtRNA1 paralogs does not affect OC43 coronavirus replication or survival of HEK293T cells. [a]** Wild-type (WT) and vt1-TKO HEK293T cell lines were infected with OC43 coronavirus, RNA purified after 24 h, then relative viral load quantified via RT-qPCR with primers for OC43 genomic RNA (gRNA) and GAPDH. **[b]** Mixed populations of wild-type and vt1-TKO HEK293T cells were seeded, infected with OC43 coronavirus ("OC43 Inf. +") or mock infected ("OC43 Inf. -"), grown under standard conditions for 48 or 72 h, then harvested. The cell lines' proportions were assessed via gDNA isolation, PCR with indicated primer sets ("Primers"; see Fig. 2a), and agarose gel electrophoresis (top). The prevalence of vt1-TKO versus WT cells ("Gene Ratio") is the ratio of "ko" and "wt" product bands, each normalized to their respective "adj" product band. Quantifications of fraction knockout cells are shown as bar graphs (bottom).

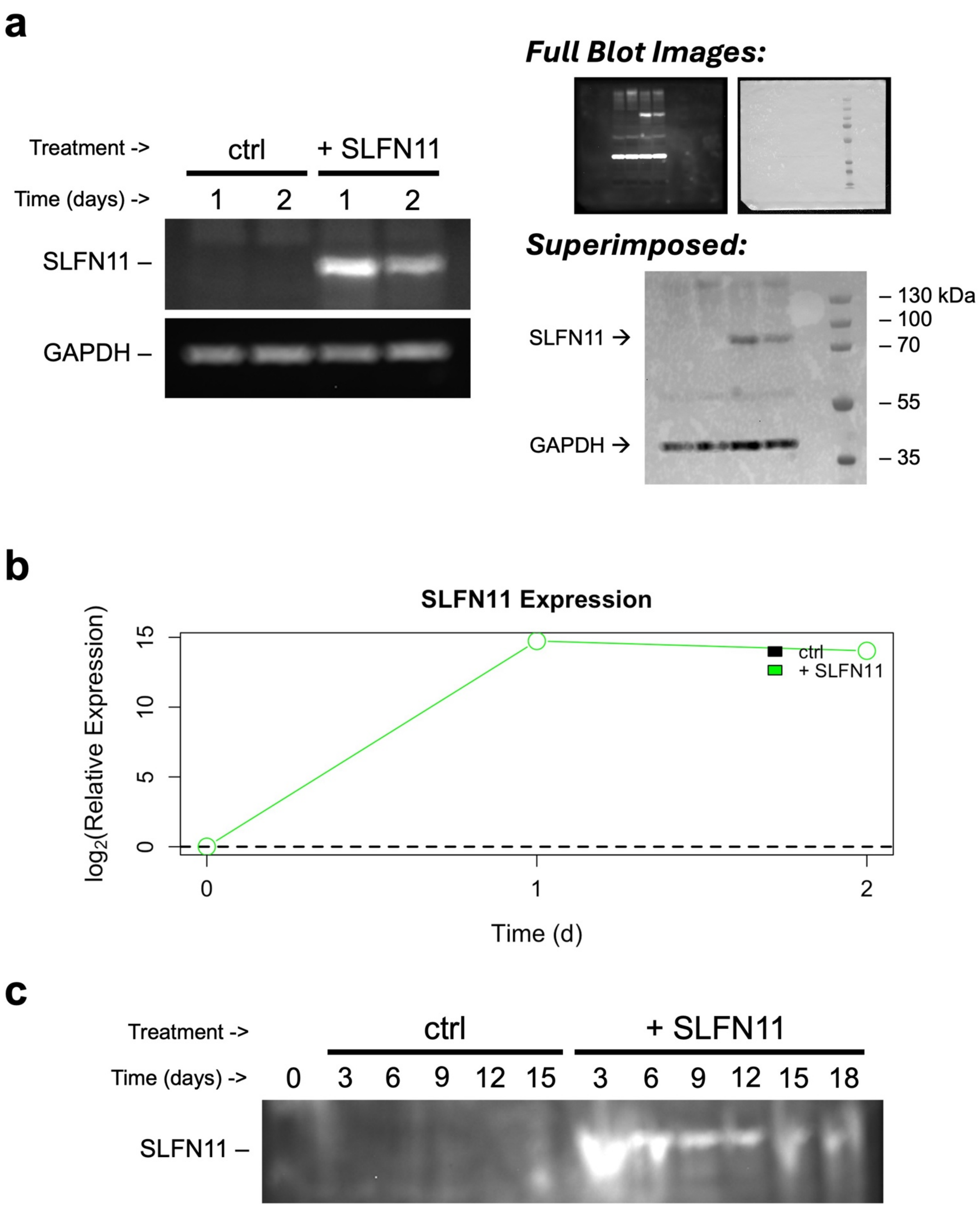


**SUPPLEMENTAL FIGURE 7. SLFN11 is successfully overexpressed in HEK293T cells. [a]** Wild-type HEK293T cells were grown under standard conditions ("ctrl") or with SLFN11 overexpression ("+ SLFN11"), sampled at 1 and 2 d post-transfection, and probed for SLFN11 expression by western blot with GAPDH loading control. SLFN11 and GAPDH bands shown (left images) are from two separate exposures of the same blot. Probe targets ("Full Blot Images", left) and protein standard ("Full Blot Images", right) were visualized with separate channels, then the blot images superimposed for molecular weight comparison (“Superimposed”). Respective band identities are indicated at left of images, and sizes for protein standard are indicated at right. **[b]** Quantification of relative SLFN11 RNA expression from HEK293T cell samples in panel-a experiment. Values were first normalized to GAPDH, then to the control (“ctrl”, dashed black line) sample on the same day. Day-0 relative expression is inferred. **[c]** Western blot of SLFN11 protein expression in mixed HEK293T cell samples from Figure 5 experiments.

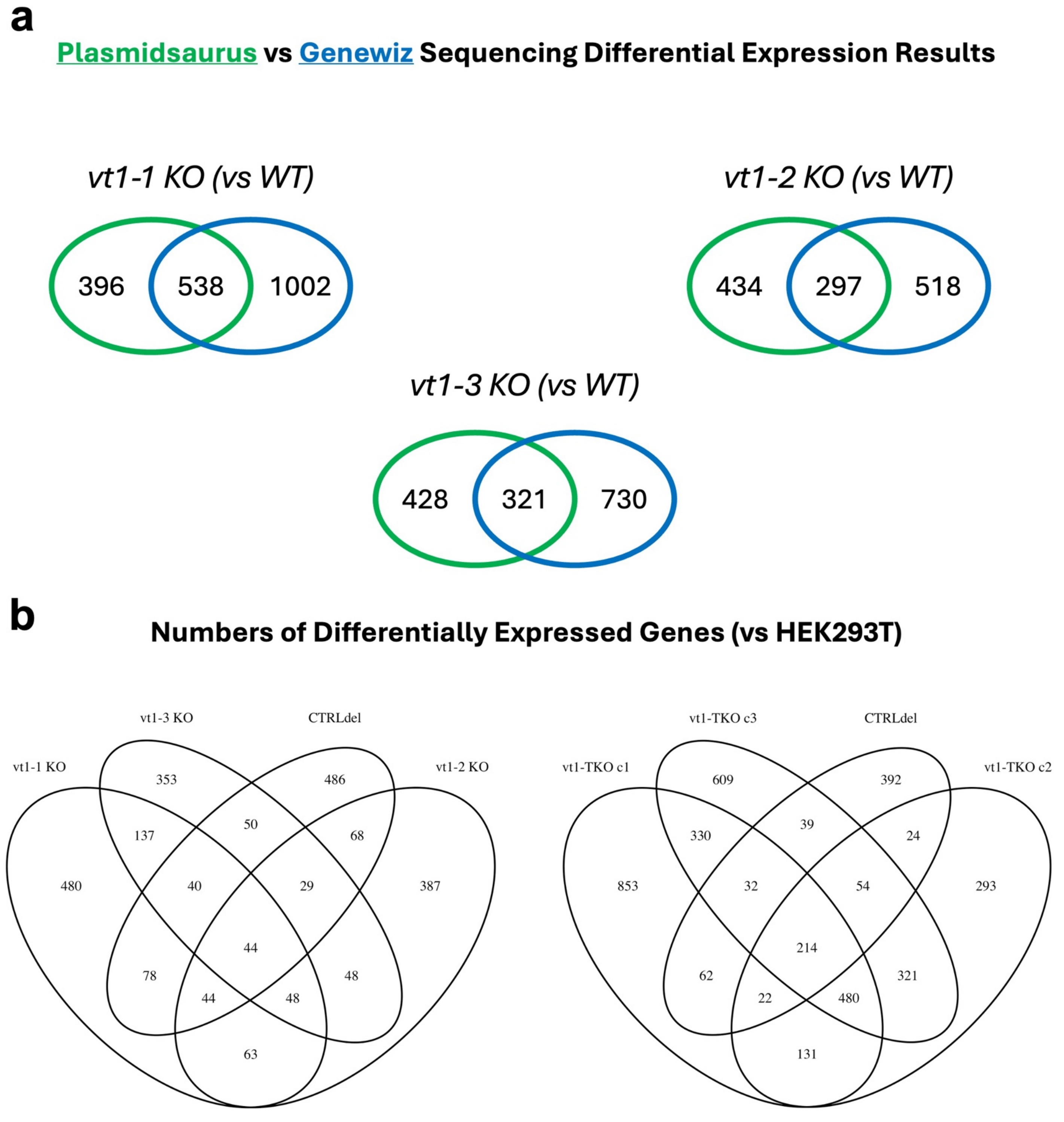


**SUPPLEMENTAL FIGURE 8. Differential expression results are significantly influenced by clonal variation and sequencing approach. [a]** Overlap between indicated RNA-sequencing methods in genes identified as differentially expressed in indicated knockout cell lines (versus wild-type; FDR < 0.05 & $\log_2|FC| > 1$). **[b]** Overlap between indicated knockout cell lines in genes identified as differentially expressed (versus wild-type; FDR < 0.05 & $\log_2|FC| > 1$) from the Plasmidsaurus sequencing set.

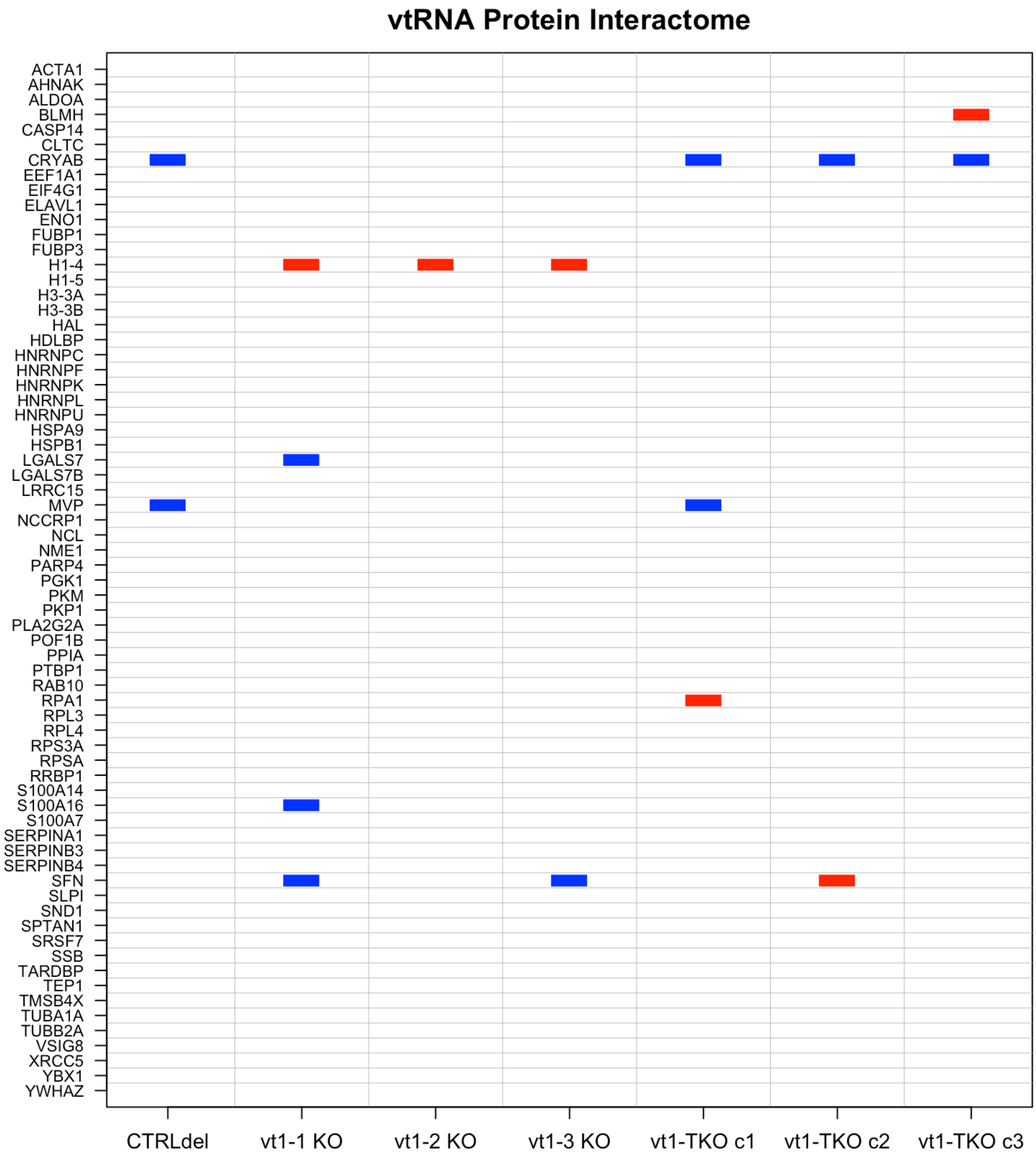


**SUPPLEMENTAL FIGURE 9. Suspected vtRNA-binding proteins are differentially expressed in HEK293T cells lacking vtRNA1 paralogs.** Suspected vtRNA-binding proteins implicated in (Stok et al. 2025) were identified in our differential expression data (Fig. 5b). Red and blue respectively correspond to down- and upregulation (FDR < 0.05 & $\log_2|FC| > 1$) of the indicated genes in the indicated cell lines.
